# Dynamic Reorganization of Developmental to Adult Genome Topology Controls the Initiation and Stabilization of the Human Muscle Stem Cell State

**DOI:** 10.64898/2026.01.23.701436

**Authors:** Matthew A. Romero, Peggie Chien, Chiara Nicoletti, Hanna L. Liliom, Gabriella Cox, Emily Skuratovsky, Kholoud Saleh, Devin Gibbs, Lily Gane, Dieu-Huong Hoang, Luca Caputo, Jimmy Massenet, Débora R. Sobreira, Pier Lorenzo Puri, April D. Pyle

## Abstract

Developmental gene expression is under tight temporal and spatial control. This regulation is imparted by tissue specific enhancers that integrate developmental signals into transcriptional responses to allow for developmental progression. Often species specific, the enhancers that regulate human muscle progenitor and stem cell gene expression are currently unknown. Here, we define the 3D chromatin organization of human muscle development and reveal key changes across the human genome that are associated with multiple layers of 3D genome reorganization during the transition from a more progenitor-like to muscle stem cell state, including a reduction of TAD numbers and an increase in CTCF binding at TAD boundaries and chromatin loops throughout developmental progression. Specifically, we found that increased CTCF occupancy at human enhancers of *PAX7* in stem cells holds enhancer-promoter (e-p) loops for timely activation of *PAX7* enhancers during early human development. These findings demonstrate that stem cell state acquisition is stabilized earlier than previously known and provide unprecedented insights into the initiation and control of the muscle stem cell state in humans.

## Introduction

Skeletal muscle possesses the remarkable ability to regenerate after an injury, largely due to its resident stem cell population referred to as satellite cells (SCs)^1^. The SC state is acquired late in skeletal muscle development requiring precise and coordinated regulation of gene expression in space and time^2^. The ability to quickly and robustly activate developmental gene expression programs is driven by the communication of *Cis*-regulator elements such as gene promoters and enhancers^3,4^. Enhancers are key elements as they govern precise spatio-temporal gene expression by binding tissue-specific transcription factors (TFs) and relaying regulatory information to their target genes across large distances. In addition to being tissue-specific, a large proportion of enhancers are species- and condition-specific. Tight regulation of the muscle stem cell state is critical for enabling the regenerative state of muscle. Despite the critical importance of muscle stem cells in muscle development and regeneration, the enhancers associated with acquisition of the SC state in human development remain unclear.

Enhancers regulate their target genes in three-dimensional space. Chromosome conformation capture (3C) techniques, such as Hi-C, have been instrumental in determining 3D chromatin topology as a key hierarchical regulator of gene expression^5^. This includes the identification of distinct features such as chromatin compartments, topologically associating domains (TADs), and chromatin loops^6,7^. To date, most studies investigating chromatin dynamics in myogenic progenitors and SCs have primarily used mouse systems^8–10^. However, a large proportion of chromatin loops and distal enhancers are species-specific, emphasizing the need to investigate chromatin dynamics in human tissues directly.

We and others have previously shown that human skeletal muscle progenitors (SMPCs) and muscle stem cells (SCs) are molecularly and functionally distinct^11,12^. How the human adult muscle SC state is established and controlled is not known. In this study we aimed to identify how the 3D genome is rewired across progenitor and stem cell states in human development and adult to determine molecular insight into when a more regenerative muscle stem cell state is obtained and regulated in human development. To this purpose, we performed a comprehensive analysis of chromatin architecture across human muscle development to adulthood. We show that during the transition from human development to the adult muscle stem cell state, myogenic gene expression is affected by regulatory changes at the level of compartments, TADs, chromatin loops, and enhancer activation and each provide key targets for locking in the muscle stem cell state. Thus the stem cell state is regulated by many critical chromatin regulatory layers. Moreover, we identified human *PAX7* super-enhancers (SEs) and enhancer stripes that become active at later human developmental timepoints signaling the initiation of control of the muscle stem cell state in humans. These findings define the 3D chromatin organization of human muscle development and reveal key changes across the human genome that coincide with multiple layers of regulation controlling the transition from early skeletal muscle progenitor to the muscle stem cell state. These findings provide key insight into the regulatory compartments important for stabilizing the muscle stem cell state that could have broad implications across both basic stem cell biology and regenerative medicine.

## Results

### Global compartment reorganization across human muscle progenitor and satellite cells

We performed Hi-C and promoter capture Hi-C (pcHi-C) analysis across key developmental stages of skeletal muscle, including early hPSC-derived SMPCs (hPSC-SMPCs), later-stage fetal-SMPCs, and adult satellite cells (SCs) (**Figure 1A, S2A,** and **S2B**). To model early human muscle development, we used *in vitro* directed differentiation to generate hPSC-SMPCs that express *PAX7*^12,13^ (**Figures S1A, S1B,** and **S1C**). Our previous work demonstrated that hPSC-SMPCs derived from this approach closely resemble early embryonic SMPCs around human weeks 7-12 in development^11^. Later-stage Fetal-SMPCs and adult SCs were isolated from *in vivo* tissues (**Figures 1A** and **S1B**). All SMPCs and SCs were freshly sorted via FACS (methods) and processed for downstream Hi-C and pcHi-C experiments. To further characterize CTCF binding and enhancer-promoter (e-p) dynamics, we performed Cleavage Under Targets and Tagmentation (CUT&Tag) for CTCF, H3K4me3, H3K4me1, H3K27ac and H3K27me3.

**Figure 1.**
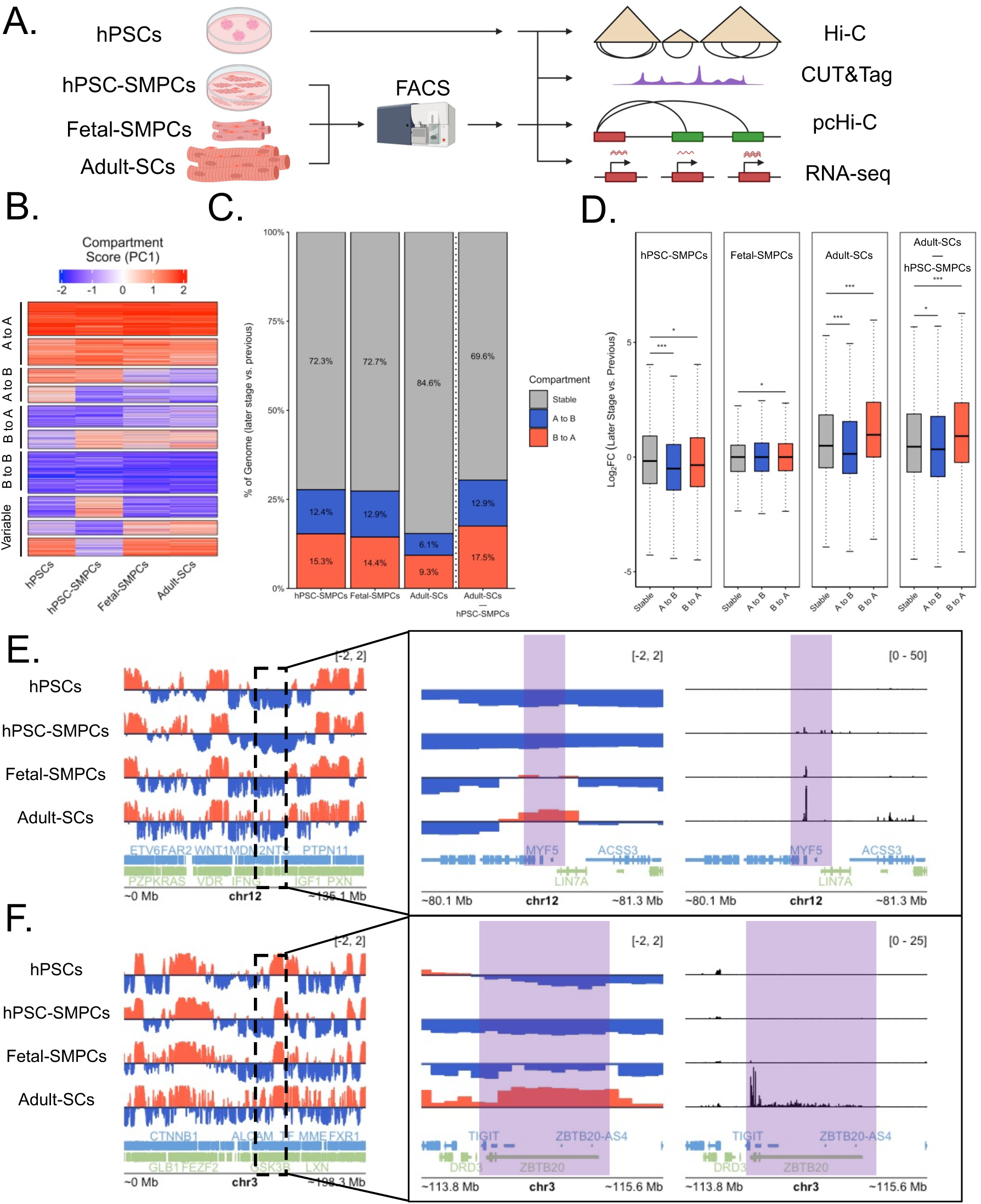
Chromatin compartmentalization across muscle progenitor development is globally reorganized in a stage-specific manner. **A)** Layout of isolation procedures and experiments conducted on hPSCs, *in-vitro* derived SMPCs from hPSCs after directed differentiation, freshly sorted fetal-SMPCs, freshly sorted adult-SCs. **B)** Heatmap displaying PC1 values used to call compartments and their dynamics across all four developmental timepoints. Y axis labels groups identified via *k*-means clustering (*k* = 6) and their compartment switching behavior at any one timepoint. **C)** Proportion of compartment switching between a later developmental timepoint and earlier stage. Note hPSC-SMPCs vs. adult-SCs. **D)** Quantification of Log2 fold change (Log2FC) in compartment identity for the regions shown in panel C. **E)** Zoomed in genome track of the *MYF5* locus depicting gradually shifting compartments corresponding to a gradual increase in *MYF5* expression (right panel in black). **F)** Genome track zoomed in on *ZBTB20* locus highlighting a sharp compartment shift in adult-SCs only, concomitant with a stark increase in gene expression (right panel in black). Mann-Whitney U test was used for significance testing. Significance is displayed as *p < 0.05, **p < 0.001, ***p < 0.0001.

To assess how large-scale chromatin compartments influence the expression of myogenic regulators, we used the first principal component (PC1) from Hi-C data, as it defines active (A) and repressive (B) compartments (**Figures S2C** and **S2D**). Using a pairwise analysis, we classified regions as stable, dynamic (switching between A and B), or highly variable (switching depending on stage) (**Figure 1B**). Pairwise comparisons between “earlier” and “later” timepoints (e.g., hPSCs vs. hPSC-SMPCs; hPSC-SMPCs vs. fetal-SMPCs) revealed that ∼30% of the genome undergoes compartment switching early in myogenesis, whereas only ∼15% is dynamic when comparing fetal-SMPCs to SCs (**Figure 1C**). We then compared gene expression profiles for genes that were in stable or dynamic compartments. We found that genes switching from B to A generally increased in expression, while those switching from A to B decreased, although these effects were less pronounced between hPSC-SMPCs and fetal-SMPCs (**Figure 1D**). When exploring specific myogenic TFs, we found that *MYF5* gradually transitioned from the B compartment in hPSCs/hPSC-SMPCs to the A compartment in fetal-SMPCs and SCs, mirroring its stepwise transcriptional activation (**Figure 1E**). To further probe how other myogenic TFs were regulated, we investigated *ZBTB20*, a zinc finger TF implicated in myogenic progression^14^. *ZBTB20* compartment switching is starker, taking place between fetal-SMPCs and SCs concomitant with a large increase in gene expression suggesting that *ZBTB20* may be involved in maturation of SCs (**Figure 1F**). Surprisingly, other canonical myogenic regulators, including *PAX7* and *MYOD*, remained in the same compartment across all stages, while *PAX3* cycles between compartments (**Figures S3A, S3B, S3C,** and **S3D**). Together, these data highlight that large-scale compartment changes are most pronounced between early and late SMPCs, with fewer changes from fetal-SMPCs to SCs. Most notable, not all myogenic regulators switch compartments until the SC stage, and some TFs don’t switch compartments at all, suggesting additional regulatory layers are at play to control the expression of myogenic TFs.

### Progressive reduction and stabilization of TADs during myogenic development

Next, we examined 3D genome architecture at the intermediate scale by identifying TADs across developmental stages. We identified 8,284 TADs in hPSCs, 7,274 in hPSC-SMPCs, 6,675 in fetal-SMPCs, and 6,371 in adult-SCs (**Figure 2A**). Consistent with previous reports, TAD size increased while TAD number decreased during myogenesis (hPSCs to adult-SCs), suggesting co-regulation of genes during development^15^ (**Figures S4A** and **S4B**). Given the role of CTCF in establishing TAD boundaries and regulating stem cell states^16–18^, we performed CUT&Tag for CTCF^19–21^. To understand if CTCF binding is differentially regulated across muscle development at TAD boundaries, we intersected CTCF peaks and TAD boundaries and calculated the proportion of TAD boundaries with CTCF bound at one, both or neither TAD boundary. We found that CTCF binding at TAD boundaries is quite dynamic across developmental timepoints (**Figure 2B**). The proportion of TAD boundaries bound by CTCF generally increased across myogenic developmental timepoints. To assess the functional impact of CTCF binding, we calculated insulation scores (IS) for boundaries with and without CTCF. At later developmental timepoints, CTCF-bound boundaries exhibited slightly weaker insulation strength than unbound boundaries (**Figure 2C**) suggesting that CTCF is aiding in the reorganization of TADs as myogenic progenitors mature^22–24^. With this in mind, we used DiffDomain^25^ to investigate TAD reorganization across our samples (**Figure 2D** and **S5**). Approximately 22-24% of TADs were reorganized, and like compartmentalization, TAD reorganization was dampened when comparing fetal-SMPCs to adult-SCs, suggesting most TAD reorganization happens in early development. We then used gene ontology (GO) analysis to understand how reorganized TADs affected developmental processes (**Figures S4C, S4D, S4E,** and **S4F**). Pairwise GO analysis revealed that TAD reorganization between hPSCs vs. hPSC-SMPCs and hPSC-SMPCs vs. fetal-SMPCs enriched for muscle development terms (**Figures S4C** and **S4D**). When comparing fetal-SMPCs vs. adult-SCs and hPSC-SMPCs vs. adult-SCs, terms associated with ECM and cell cycle regulation were enriched, respectively (**Figures S4E** and **S4F**). While the amount of TAD reorganization is reduced at later timepoints, the proportion of TAD reorganization types was consistent when comparing pairwise developmental timepoints (**Figure 2E**). An example of TAD reorganization happening in early development then staying stable across myogenic development is the *HOXA* cluster (**Figure 2F**). This reorganization coincides with increased *HOXA* gene expression, highlighting the need for precise spatial regulation of this cluster during muscle development^26^. Similar to compartments, however, many myogenic TFs do not undergo a rewiring of TADs (**Figures S7A** and **S7B**), suggesting their regulation is mediated at finer scales, such as chromatin loops or enhancer–promoter interactions within insulating neighborhoods.

**Figure 2.**
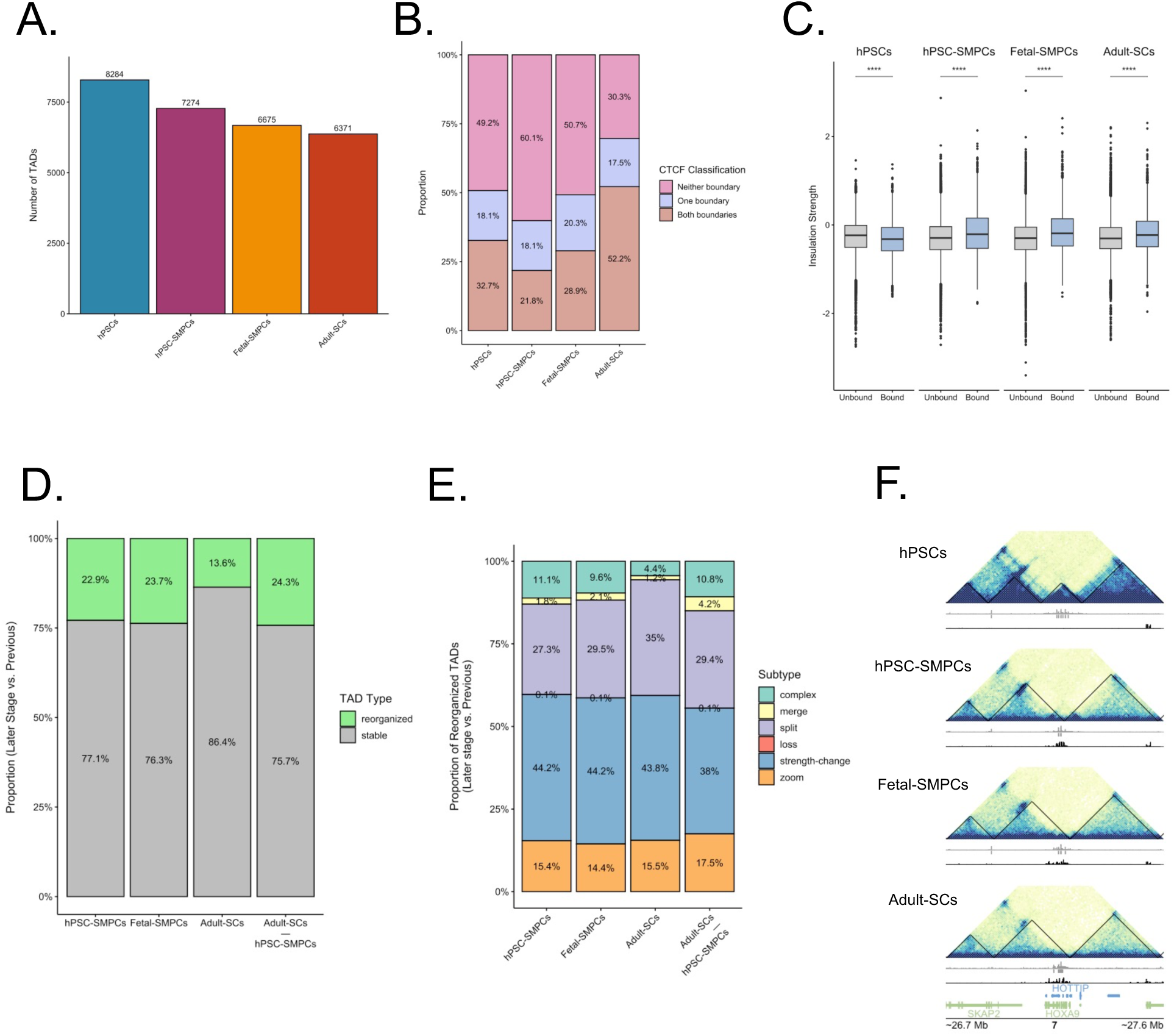
Global TAD reorganization occurs across muscle development, but only at a subset of myogenic regulators. **A)** Total number of TADs per developmental condition. **B)** Proportion of TADs that are bound by the boundary protein CTCF. CTCF binding assessed via CUT&Tag. **C)** TAD border strength as measured by the Insulation Score and its relationship with CTCF binding. **D)** Proportion of TADs that undergo reorganization when comparing earlier developmental timepoint with the subsequent developmental timepoint. Note the comparison of hPSC-SMPCs vs. adult-SCs. **E)** Quantification of TAD reorganization events from analysis with DiffDomain when comparing earlier developmental timepoint with the subsequent developmental timepoint. Note the comparison of hPSC-SMPCs vs. adult-SCs. **F)** Pyramid plot around the HOXA loci depicting TAD reorganization after specification of hPSC-SMPCs and this stays consistent across the remaining developmental timepoints. Mann-Whitney U test was used for significance testing. Significance is displayed as *p < 0.05, **p < 0.001, ***p < 0.0001.

### Chromatin loops are dynamically rewired across development

To better understand fine-scale chromatin folding, we used our Hi-C data to investigate the dynamics of chromatin loops across our four developmental timepoints^27^. We found 11,300 loops in hPSCs, 12,007 in hPSC-SMPCs, 9,166 in fetal-SMPCs, and 8,751 in adult-SCs (**Figure S8A**). Loop overlap analysis revealed that the majority of loops are stage-specific, with hPSCs and hPSC-SMPCs containing the largest numbers of unique loops (**Figures 3A** and **S8B**). In contrast, fetal-SMPCs and adult-SCs shared a similar number of unique loops. The largest proportion of loop overlaps occurred between hPSC-SMPCs and fetal-SMPCs, followed by loops shared among all myogenic cells and those shared between fetal-SMPCs and adult-SCs. Following our TAD analysis, we examined CTCF occupancy at loop anchors, classifying loops as bound at one, both, or neither anchor. Consistent with TAD-level results, the proportion of CTCF-bound loops was highest in adult-SCs (**Figure 3B**). Loop strength, measured via aggregated peak analysis (APA), also peaked in adult-SCs, consistent with patterns seen during terminal differentiation (**Figure 3C**). We then intersected CTCF peaks with loop anchors and analyzed loop strength via APA. We show a similar trend in that CTCF anchored loops were strongest in adult-SCs suggesting CTCF is also involved in either initiating or reinforcing stem cell states at the level of chromatin looping (**Figure 3D**). CTCF plays a complex regulatory role and CTCF anchored loops can either help promote or repress gene expression^28^. To better capture and determine CTCF’s role in gene expression across myogenic development, we calculated the proportion of differentially expressed genes (DEGs) with CTCF bound to their promoter. We found that the proportion of CTCF at the promoters of DEGs is seen at both up and downregulated genes in hPSC-SMPCs compared to hPSCs (**Figure 3E**). The proportion of DEGs with CTCF at their promoter is somewhat blunted when comparing hPSC-SMPCs with fetal-SMPCs; however, when comparing fetal-SMPCs and adult-SCs, CTCF was largely found at promoters of downregulated genes. We then filtered our analyses for GO analysis based on the proportion of CTCF bound to up- or downregulated genes. GO analysis of CTCF bound downregulated genes in adult-SCs identified enrichment for mitosis related process when compared to fetal- and hPSC-SMPCs (**Figures S8D** and **S8E**). Downregulated genes in Fetal-SMPCs compared to hPSC-SMPCs enriched for metabolic processes (**Figure S8F)**. In hPSC-SMPCs, we used GO analysis to investigate CTCF bound upregulated genes and found an enrichment of muscle associated processes suggesting that CTCF may aid in the upregulation of muscle-specific genes in the early stages of muscle development (**Figure S8G**).

**Figure 3.**
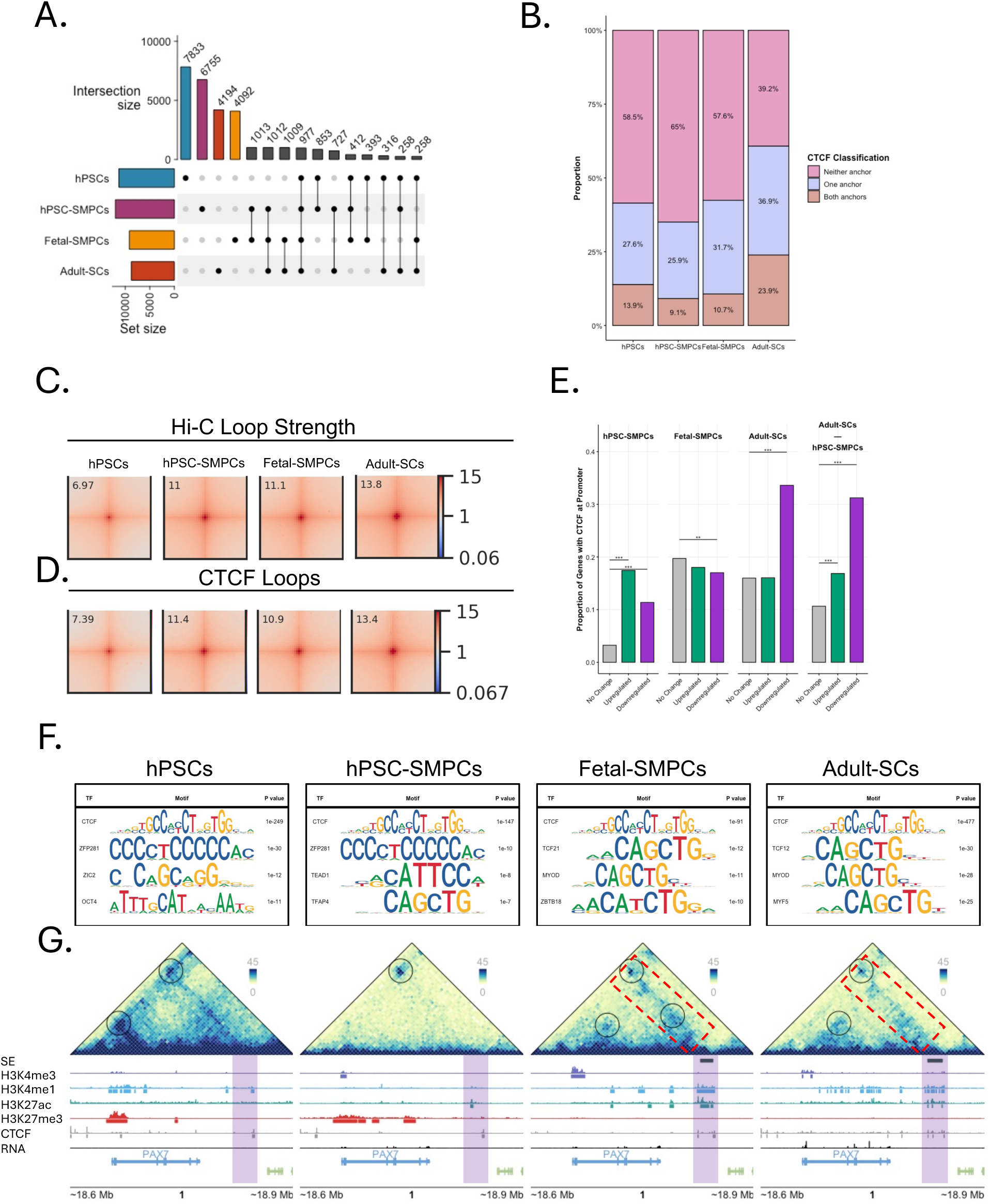
Myogenic loop dynamics reveal stage specific regulatory control. **A)** Upset plot depicting amount of unique and overlapping loops between conditions. Bars are depicted in descending order. **B)** Proportion of CTCF bound at chromatin loops as called by Hi-C. **C)** chromatin loop strengths quantified via APA across all loops and **D)** CTCF bound loops. **E)** Quantification of proportion of DEGs with CTCF binding at their promoters. Comparisons are made between earlier vs. later developmental timepoint. Note hPSC-SMPCs vs. adult-SCs. **F)** Motifs enriched at H3K4me1 chromatin loops across all four conditions using HOMER. **G)** Pyramid plot around the *PAX7* locus depicting chromatin loops between the *PAX7* promoter and putative enhancers. Red dashed box depicts an enhancer stripe called via the Stripenn tool. Purple opaque vertical bars are highlighting the putative enhancer regions for *PAX7*. Mann-Whitney U test was used for significance testing. Significance is displayed as *p < 0.05, **p < 0.001, ***p < 0.0001.

### Enhancer–promoter loops and transcription factor motif enrichment

Enhancers play a critical role in temporal and cell type–specific transcriptional control^29^. To explore all enhancer anchored loops, we intersected H3K4me1 CUT&Tag peaks with our chromatin loop anchors from Hi-C data. We scanned for motifs enriched in H3K4me1 peaks within loop anchors using HOMER and identified possible transcription factors that may initiate or otherwise regulate loops at each specific developmental timepoint (**Figure 3F**). We found CTCF motifs enriched across all four timepoints as well as lineage determining TFs unique to each stage that include *OCT4* (hPSCs), *TEAD1* (hPSC-SMPCs), *MYOD* (fetal-SMPCs and adult-SCs) and *MYF5* (adult-SCs). Given previous reports that CTCF interacts with lineage determining TFs to regulate gene expression, these data suggest that these TFs may be interacting with CTCF to hone and establish developmentally specific loops across the genome^30,31^.

### *PAX7* regulatory architecture

*PAX7* plays a very important role in myogenesis and is required for SC specification and maintenance in both development and adulthood^2,32^. Investigating the regulatory sequences that control *PAX7* expression is difficult to study due to the fact that *PAX7* expression is abolished during culturing. Leveraging our freshly isolated hPSC-SMPCs, fetal-SMPCs and adult-SCs, we integrated Hi-C loop maps with enhancer-associated histone modifications from CUT&Tag to identify candidate *PAX7* regulatory elements. We identified multiple intergenic regions interacting with the *PAX7* promoter, which displayed both looping interactions and enhancer-associated chromatin marks (**Figure 3G**). These loops were not unique to one stage and were largely maintained across all four samples. The enhancer modifications, however, became enriched in hPSC-SMPCs and were highest in adult-SCs. The inverse was seen for the repressive chromatin mark H3K27me3. Fetal-SMPCs and adult-SCs show a depletion of H3K27me3 at the *PAX7* promoter, while H3K27me3 remains in both hPSCs and hPSC-SMPCs. Similarly, the enrichment of CTCF at enhancer regions were highest in adult-SCs. These data suggest that CTCF may be stabilizing long-range regulatory contacts between the *PAX7* promoter and distal enhancers^33^.

### Super enhancers and enhancer stripes at *PAX7*

Super enhancers (SEs) and enhancer stripes are frequently associated with lineage-defining TFs and highly expressed genes, respectively^34–36^. Using ROSE (for SEs) and Stripenn (for enhancer stripes) algorithms, we found that both features became enriched at *PAX7* in late fetal-SMPCs and adult-SCs, concomitant with the highest expression level of *PAX7* (**Figure 3G**). These results suggest that chromatin loops at the *PAX7* locus are established early in development and remain, but that enhancers controlling the stem cell state do not become activated until late fetal myogenic development. Importantly this work showed that stabilization of the stem cell state is acquired earlier than previously anticipated in late fetal development.

### Promoter capture Hi-C identifies stage-specific enhancers

While our Hi-C data highlighted candidate regulatory regions for *PAX7*, its resolution at gene promoters limits the ability to assign e-p interactions. To enrich for promoter-anchored interactions, we performed pcHi-C across all four developmental stages^37^. We found 136,029 loops in hPSCs, 149,342 in hPSC-SMPCs, 141,035 in fetal-SMPCs, and 150,620 in adult-SCs, respectively (**Figure S9A**). Similar to Hi-C data, hPSCs contained the largest proportion of unique loops (**Figure 4A**). However, unlike Hi-C, pcHi-C revealed that loop overlaps are highest in all four conditions, suggesting a shared e-p network across multiple stem cell states. We next sought to investigate all activated enhancer-promoter loops (e-p loops) by intersecting H3K27ac CUT&Tag peaks with pc-HiC loop anchors across all four conditions. We normalized e-p loop counts across each gene and identified five distinct temporal patterns across conditions (**Figure 4B**). These loop count patterns spanned e-p loops that were enriched in hPSC-SMPCs and fetal-SMPCs (pattern 1), gradually decreased as development proceeds (pattern 2), enriched in hPSCs (pattern 3), gradually increased as development proceeds (pattern 4) and primarily enriched in adult-SCs (pattern 5). We then filtered our intersected dataset for DEGs and conducted GO analysis. We found distinct processes enriched in each pattern (**Figure S9B)**, for example, pattern 4 enriched in muscle development DEGs, while pattern 5 enriched for ECM and collagen related DEGs. This led us to ask if e-p loops were increasing, decreasing or otherwise being lost. To assess dynamic changes, we compared conditions pairwise to look at all genes with e-p loops and categorized them as absent, constant, increased in number, or decreased in number. Notably, e-p loops largely increased or were absent when comparing conditions (**Figure 4C**). However, we did note that at later developmental timepoints (fetal-SMPCs vs. adult-SCs), a larger proportion of e-p loop counts were decreasing across genes. This suggests that e-p loops play a fine-tuning role in gene expression, by either increasing or decreasing e-p loops, at later developmental timepoints whereas earlier developmental timepoints show more of a reliance on loop addition or total loss of loops. We then quantified the impact of gained e-p loops on gene expression and found a concomitant increase in expression for genes that gained e-p loops for fetal-SMPCs and adult-SCs pointing to the importance of adding activated enhancer loops at later stages of muscle development (**Figure 4D**). Given our identification of SEs at putative *PAX7* regulatory sites, we sought to explore the impact of SEs on gene expression genome-wide. We compared the expression of genes that were anchored by either SE loops or H3K27ac loops. We found that SE anchored e-p loops have higher expression compared to H3K27ac anchored e-p loops, but notably only after specification of hPSC-SMPCs and later developmental timepoints (**Figure 4E**). Motif enrichment analysis of SE loops revealed developmental stage–specific transcription factors, including *ZIC3*, *FOXA1*, *MYOD* and *PAX7* (**Figure 4F**). These data suggest that SEs may have the ability to regulate myogenic cell states throughout muscle development, driven by TFs such as *MYOD* and *PAX7*^8^.

**Figure 4.**
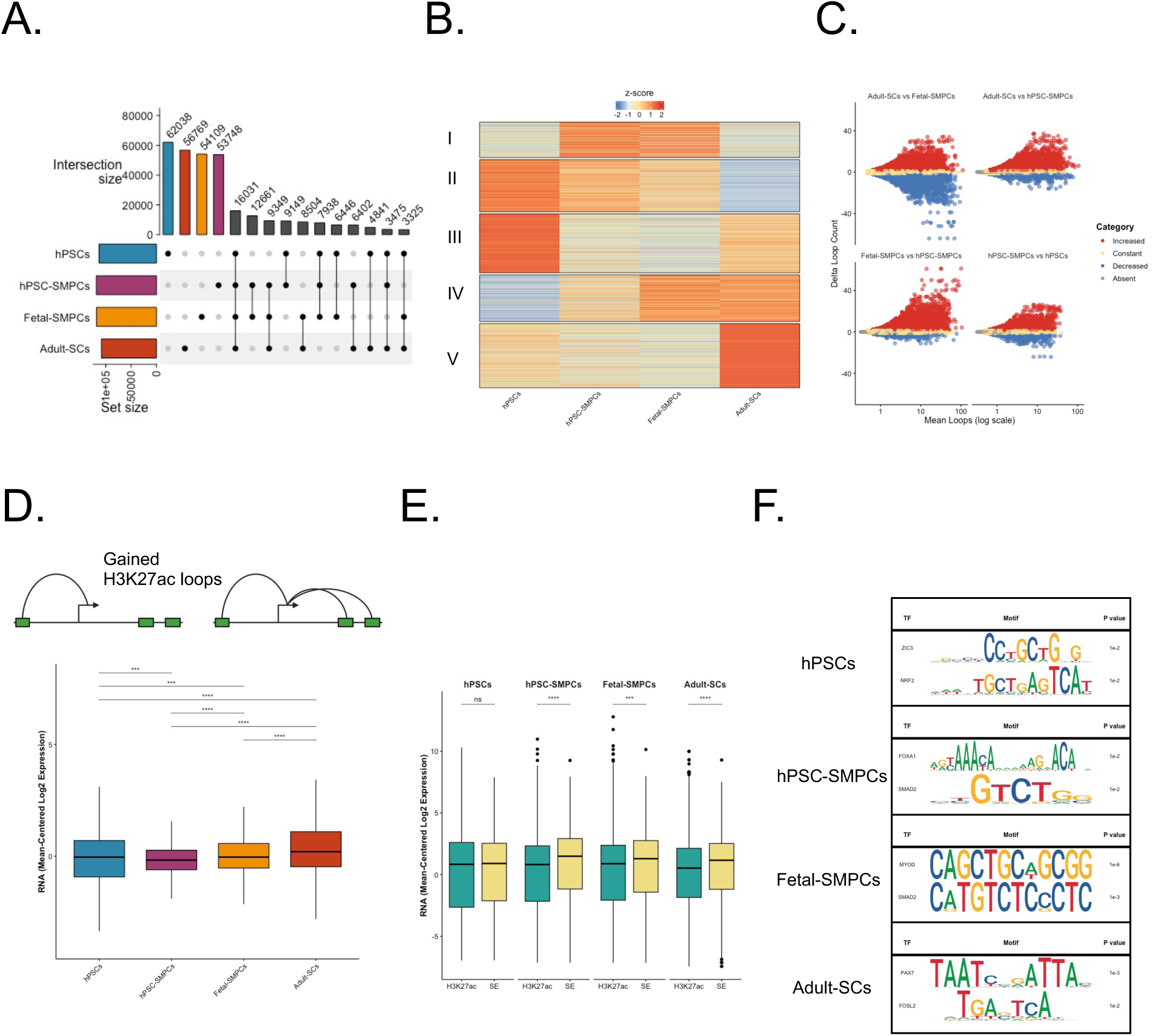
Chromatin loops via pcHi-C uncover enhancer usage across myogenic progenitors. **A)** Upset plot depicting amount of unique and overlapping loops between conditions. Bars are depicted in descending order. **B)** Clustering (*k*-means, *k* = 5) of H3K27ac anchored pcHi-C (enhancer-promoter, e-p) loops and their regulation across all conditions. **C)** Volcano plots depicting genes with e-p loops across all four conditions that are constant, gained (increased), lost (decreased), or have zero loops. **D)** Quantification of gene expression for genes that gained H3K27ac loops. **E)** Comparison of gene expression between H3K27ac anchored e-p loops and SE anchored e-p loops. **F)** Motifs enrichment found at SEs across all four conditions. Mann-Whitney U test was used for significance testing. Significance is displayed as *p < 0.05, **p < 0.001, ***p < 0.0001, ****p < 0.00001.

### Identification of *PAX7* enhancers in human skeletal muscle progenitors

*PAX7* is essential for SC specification and quiescence, but the enhancers that control *PAX7’s* expression in human muscle progenitors and SCs is unknown. Our Hi-C analysis identified one e-p loop at *PAX7* (**Figure 3G**), yet genes are often regulated by multiple, temporally controlled enhancers^38^. To comprehensively map *PAX7* regulation, we integrated pcHi-C data with CUT&Tag profiles for histone modifications, H3K4me3, H3K4me1, H3K27ac, H3K27me3 and CTCF across all four of our conditions. *PAX7* promoter-anchored loops were present in hPSCs, however, they tend to increase in number across all myogenic stem and progenitors (**Figure 5A**). A prominent feature in hPSCs and hPSC-SMPCs, but absent in fetal-SMPCs and adult-SCs, was the H3K27me3 deposition at the *PAX7* promoter, indicative of transcriptional repression, particularly in hPSCs. Downstream of the *PAX7* promoter, hPSC-SMPCs began establishing loops seen in fetal-SMPCs and adult-SCs. However, these loops initially lacked strong H3K4me1 and H3K27ac activation marks. These data suggest that there is an interplay between the establishment of e-p loops early in development and the activation of enhancers concomitant with removal of repressive chromatin marks at later developmental timepoints. Moreover, in both fetal-SMPCs and adult-SCs, CTCF occupancy at *PAX7* enhancers increased, suggesting the establishment of longer held e-p loops (**Figure 5B**). We also noted the activation of an SE at a subset of downstream *PAX7* e-p loops exclusively in fetal-SMPCs and adult-SCs. Taken together, these data highlight that multiple processes coalesce at enhancers to regulate *PAX7* expression and there is a level of both enhancer switching and enhancer stability as development proceeds toward SC maturation.

**Figure 5.**
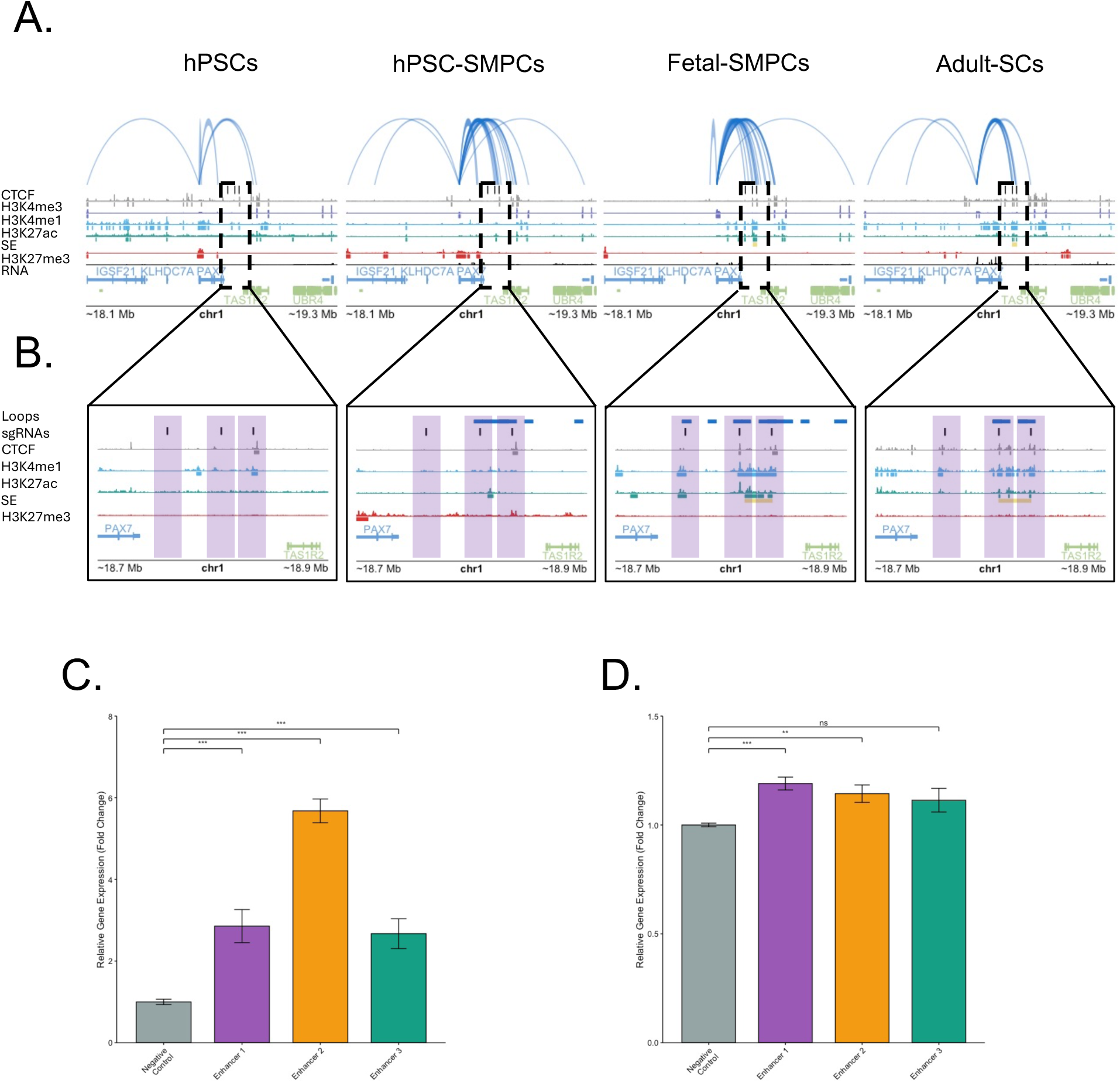
Activating *PAX7* enhancers upregulates PAX7 expression *in-vitro*. **A)** Genome track around the *PAX7* locus identifying chromatin loops anchored at the *PAX7* promoter. Genome tracks include tracks for CTCF (Grey), H3K4me3 (purple), H3K4me1 (blue), H3K27ac (green), SEs (yellow), H3K27me3 (red), and RNA (black). **B)** Zoomed in view of the putative enhancers regulating *PAX7*. Vertical black bars within vertical purple opaque bars depict candidate enhancers that are used for subsequent **C)** luciferase and **D)** CRISPR/dCas9-VPR experiments, respectively. Enhancers 1, 2, and 3 are named based on 5’ to 3’ distance from the *PAX7* gene. **C)** and **D)** Data are represented as mean +/- SD; n = 3. Statistical significance tested with two-tailed Student’s *t*-test. Significance is displayed as *p < 0.05, **p < 0.001, ***p < 0.0001, ****p < 0.00001.

### Functional validation of *PAX7* enhancers

To functionality validate candidate *PAX7* enhancers, we selected three regions for downstream luciferase reporter assays (**Figure 5B**). Each region was individually cloned into a minimal promoter-driven luciferase vector, and a genomic site lacking enhancer marks was cloned as a negative control. Transfection of these constructs into primary fetal-SMPCs showed that all three regions significantly increased luciferase activity relative to the negative control, indicating that these regions function as enhancers (**Figure 5C**). To test whether these putative enhancers could upregulate *PAX7* expression in a native context, we designed gRNAs tiling each of the three enhancer regions. Primary fetal-SMPCs were separately transfected with each gRNA tiling set together with the activating dCas9-VPR system. Enhancer 1 and Enhancer 2 each induced ∼20% increases in *PAX7* expression, whereas Enhancer 3 had no significant effect (**Figure 5D**). These findings suggest that specific *PAX7* enhancers could be used as a tool to maintain *PAX7* expression *in vitro,* offering a potential strategy to sustain *PAX7* expression and facilitate *in vitro* SC studies while maintening the stem cell state.

## Discussion

The role chromatin plays in transcriptional regulation in SMPCs and SCs in human muscle is difficult to study and has largely relied on findings in other mammalian systems, such as mouse, and *in-vitro* directed differentiation strategies using human PSCs and induced PSCs^8,9,39^. While these investigations have extended our understanding of conserved biological mechanisms, many aspects of enhancer regulation and chromatin topology are species specific^40,41^ and could be developmentally regulated. To minimize this knowledge gap, we profiled chromatin architecture and enhancer regulation across four developmentally distinct timepoints and identified that chromatin imparts transcriptional regulation across various scales, ultimately providing evidence that *PAX7* expression can be maintained *in-vitro.* These data will help design tools targeting stage-specific enhancers improving our ability to study SCs *in-vitro*; possibly by increasing our ability to maintain SC stemness by regulating *PAX7* expression and thus preventing differentiation. In parallel generation of hPSC reporter lines that read out different enhancer states could be used to facilitate screens that generate more adult like SC states in directed differentiation cultures.

At the largest scale, we found that chromatin compartmentalization is dynamic throughout muscle development but is dampened as cells “get closer” to fully mature adult-SCs. Surprising to us, however, many myogenic TFs do not switch compartments but are pre-emptively in the active A compartment. This pre-emptive compartmentalization could, in part, be due to the fact that SMPCs are actively proliferating during development and quiescent SCs are isolated via FACS, which results in cell activation^42^. However, our reports align with previous mouse studies that utilized formaldehyde fixation to preserve quiescence and reported similar myogenic TFs including *PAX7*, *PAX3*, *MYF5*, are in the A compartment in quiescent SCs and are maintained in compartment A throughout differentiation^9^. This suggests that TFs that regulate myogenic identity are held in the active compartment and rely on fine scale regulation for their expression, including e-p loop activation/addition.

We also showed a large amount of TAD reorganization across our timepoints, specifically around the HOXA gene cluster. A large proportion of TAD reorganization was a change in the boundary strength of these TADs, suggesting that interaction strength, either higher or lower, also plays a significant role in regulating gene expression^43^. Taken together with the finding that many myogenic TFs don’t show TAD reorganization, this would fall in line with a previous study identifying that many developmental genes are housed in single gene TADs and are quite stable across differentiation^44,45^. Our data recapitulate these findings as major myogenic regulators such as *PAX7*, *MYF5*, and *MYOD* do not reorganize their TADs. These data support that TADs are large scale regulators of gene expression that work to mitigate large-scale changes that would result in lethal or disease-oriented gene expression, such as seen in limb malformations and certain cases of cancers, including rhabdomyosarcoma^46,47^.

The most dynamic regulators across our samples were chromatin loops. It has been previously reported that chromatin loops strengthen across terminal differentiation^9^. Our data shows that this is not just involved with terminal differentiation, but also as cells move along a developmental trajectory, i.e., SMPCs moving toward SCs. We also highlight that CTCF may be playing a large role in downregulating gene expression as SMPCs mature into SCs, adding to the complex role that CTCF plays in gene regulation. Using pcHi-C, we were also able to identify enhancers that are active at specific developmental timepoints across myogenic progenitors. Specifically, we evaluated the enhancers that control human *PAX7* expression. *PAX7* is crucial for SC specification and maintenance, and its expression is abolished shortly after culturing. Using CRISPRa to activate our candidate enhancers, we show the ability to maintain *PAX7* expression. Understanding how *PAX7* expression is regulated will allow us to develop tools to modulate its expression in culture, and approaches to expand stem cells in culture while maintaining stemness and preventing differentiation which would be the holy grail for using muscle stem cells in translational applications.

Our study provides an overarching view of chromatin regulation across myogenic progenitor and adult muscle stem cell states and serves as a resource of *in-vivo* chromatin topology and enhancer usage at all levels. Taken together, these results suggest that chromatin regulation in human myogenesis follows a hierarchical model: broad-scale compartment organization sets a permissive or repressive chromatin state for myogenic TF loci early, TAD stability safeguards key developmental genes, while dynamic looping—especially enhancer-promoter contacts—fine-tunes transcriptional output during progression toward the SC state. We were able to identify three novel human enhancers that regulate human *PAX7* expression that are active in both fetal-SMPCs and adult-SCs demonstrating stabilization of the stem cell state occurs earlier than previously recognized. Importantly we showed that enhancer usage and stability changes as progenitors mature to adult muscle SCs. This knowledge could be used to activate specific enhancers in progenitors to readily promote transition to adult-SCs generating more mature and regenerative SC states in culture.

## Methods

### hPSC lines, directed differentiation, and cell culture

All human pluripotent stem cell work was approved by ESCRO. hPSCs were grown and maintained on Matrigel coated plates in mTESR medium (Stem Cell Technologies) containing 0.5% penicillin/streptomycin (p/s, Hyclone) and differentiation of hPSCs into SMPCs was performed as previously described^11,12^. hPSC lines were dissociated in TrypLE and plated at 275,000–475,000 cells per well of a six-well plate in mTESR containing 10 µM ROCK inhibitor for 1 day to allow for recovery. hPSCs were then treated with 10 µM CHIR99021 (Tocris, 4423) in E6 medium for 2–3 days. E6 medium was used for and additional 7–10 days. StemPro-34 medium containing 10% StemPro-34 supplement, 5 ng/ml basic fibroblast growth factor (bFGF), 2 mM L-glutamine, 0.45 mM monothioglycerol, 11 μg/ml transferrin and 0.5% p/s was used for 6–8 days. E6 medium was then added to cells for 3 days supplemented with 10 ng/ml IGF-1 for 7–10. N2 medium (1% N2 supplement, 1% insulin-transferrin-selenium (ITS, Gibco), 10 ng ml–1 IGF-1 and 0.5% penicillin-streptomycin) was added to cells for 5–7 days, and N2 medium containing TGFβi (SB431542; 3 μM) was added for 5–7 days prior to hPSC-SMPC isolation.

### Preparation of hPSC-SMPCs, fetal-SMPCs and adult-SCs

At the end of directed differentiation, hPSC-SMPCs were prepared for FACS by dissociation in TrypLE, as previously described (REFs). Cells were filtered through 100-µm filters, washed in PBS and resuspended in FACS buffer (2% FBS in PBS). HNK1^-^ERBB3^+^NGFR^+^ hPSC-SMPCs were isolated via FACs on a BD FACSAria II. Fluorescence minus one (FMO) controls were used for all gating controls. DAPI staining was used to exclude dead cells.

Fetal tissues were IRB exempt and obtained from ABR (described in table XXX). Fetal muscles (week 17-20) were dissected and washed with wash buffer and then mechanically dissociated into small pieces at RT in digestion buffer consisting of wash buffer supplemented with 2 mg/ml of Collagenase II,1 mg/ml of Dispase II and 25 µg/ml of DNase I. Chopped tissues were further incubated in digestion buffer on a shaker at 37 deg. C for 30 minutes with intermittent trituration. Digestion was stopped by adding extra FACS buffer. Digested tissues were filtered through 100 μm cell strainers and spun down. Cell pellets were resuspended in FACS buffer, filtered through 70 µm cell strainers, spun down and resuspended again in FACS buffer at 1×10^6^ cells per ml. Fetal-SMPCs were sorted on CD31^-^CD45^-^CD235a^-^NCAM^+^CD82^+^ via FACs on BD FACSAria II sorters. DAPI staining was used to exclude dead cells.

Human adult quadriceps muscle was donated irregularly from donor cadavers from NDRI. Adult quadriceps muscle was dissected and washed with wash buffer and then mechanically chopped into smaller pieces at RT in primary digestion buffer consisting of wash buffer supplemented with 2 mg/ml of Collagenase II. Chopped tissues were further incubated in primary digestion buffer on a shaker at 37 deg C for 20 minutes with intermittent trituration. Primary digestion was stopped by adding extra wash buffer and tissues were spun down. Next, supernatant was removed, and tissues were resuspended in secondary digestion buffer consisting of wash buffer supplemented with 7 mg/ml of Collagenase D, 1.5 mg/ml of Dispase II and 25 µg/ml of DNase I. Tissues were further digested on a shaker at 37 deg C for 20 minutes with intermittent trituration. Secondary digestion was stopped by adding extra FACS buffer. Digested tissues were filtered through 100 µm cell strainers and spun. Cell pellets were resuspended in FACS buffer, filtered through 70 µm cell strainers, spun down and resuspended again in FACS buffer at 1×10^6^ cells per ml. Adult-SCs were sorted on CD31^-^CD45^-^CD235a^-^NCAM^+^CD82^+^ via FACs on BD FACSAria II sorters. DAPI staining was used to exclude dead cells. Our current study only describes the changes seen across human muscle development and due to limitations in sample availability we were required to use *in-vitro* derived hPSC-SMPCs to model earlier stages of human embryonic muscle development. Although our previous studies determined that hPSC-SMPCs align with early embryonic progenitors^11^ our model may not fully recapitulate chromatin architecture changes happening *in-vivo*. Furthermore, although our samples were processed immediately after sorting, the procedure required for the isolation of SMPCs and SCs via FACS activates SCs gene expression within a few hours. This means that both SMPCs and SCs were not truly in the quiescent state. Therefore, the role chromatin architecture plays in quiescence will need better tools and isolation strategies to evaluate genome studies as performed here in quiescent SCs. Given the rarity of these populations and donations of tissue, certain samples had to be pooled in order to conduct our analyses. Recent improvements in single- and low-cell input protocols will be needed to further shed light on how chromatin topology influences gene expression in single cell populations.

### Hi-C and pcHi-C

Two biological replicates were used for hPSCs, hPSC-SMPCs, and fetal-SMPCs. Each replicate consisted of a cell count between 500,000-1,000,000 cells. Adult-SCs are an extremely rare population. As such, SCs were isolated via FACs from multiple donors and pooled together, approximately 150,000 cells, prior to Hi-C experiments using ARIMA Genomics Hi-C^+^ kit. Sorted cells were fixed in 2% formaldehyde, quenched according to manufacturer’s instructions, spun down and the remaining pellet was stored at −80 deg C until further processing. Hi-C was performed according to manufacturer’s protocol. Between 1-1.5 µg of DNA was sonicated on a Covaris M220 (DutyFactor=20%, time=150s, Incident Power=50W) to generate ∼400bp DNA fragments. Biotinylated fragments were purified using streptavidin beads. 200 ng of sonicated and purified DNA was used for library preparation using ARIMA’s indexed primers. A fraction of Hi-C library was used for sequencing. Paired-end reads of 150bp were obtained using Illumina NovaSeq 6000. The remaining fraction of Hi-C library was used as input for pcHi-C using ARIMA Genomics protocol using a custom panel of human promoters. After promoter capture, libraries were amplified according to the manufacturer’s instructions. The subsequent pcHi-C libraries were sequenced (paired-end, 150bp) using an Illumina NovaSeq 6000.

### Data processing for Hi-C

Raw sequencing files were processed using Juicer v1.6 with genome assembly hg38 and ARIMA’s restriction sites^53^. FASTQ’s of biological replicates were merged prior to running juicer. The resulting hic files were converted to cool files using HiCExplorer’s hicConvertFormat tool^27^. Subsequent cool files were read depth normalized and KR corrected. Compartments were called using dcHiC at 100kb resolution and H3K27ac CUT&Tag peaks were used to properly assign the PC1 results^49^. TADs and chromatin loops were called using HiCExplorer at 25kb and 5kb resolution, respectively. DiffDomain was used to call reorganized TADs from the results of TAD calls from HiCExplorer. Aggreate Peak Analysis (APA) was calculated via Open2C’s coolpup.py analysis tool^54^. Enhancer stripes were called via the Stripenn tool with standard options^34^. Visualization for Hi-C data was completed using Plotgardener^51^.

### Data processing for pcHi-C

Raw sequencing files were processed using ARIMA’s bioinformatics pipeline for pcHi-C (Arima-CHiC-v1.3). The HiCUP pipeline was used to map fragments using bowtie2 using the hg38 assembly^55^. BAM files were then converted to CHiCAGO input files and subsequently processed using CHiCAGO to obtain interaction confidence scores^52^. Only scores above or equal to 5 were used for downstream analysis. Significant interactions were used to integrate with enhancers generated from CUT&Tag experiments. Visualization was completed using Plotgardener^51^.

### CUT&Tag

CUT&tag was performed as described previously^19,20^. Freshly sorted cells were isolated via FACS as described above and immobilized on Concanavalin A-coated paramagnetic beads and permeabilized with digitonin. Cells were then incubated overnight with antibodies targeting histone modifications (H3K4me3, H3K4me1, H3K27ac, H3K27me3), CTCF, or negative control (IgG). Cells were then washed to remove unbound antibody and treated with pA-Tn5 (Epicypher) for 1hr. Tn5 was then activated via the addition of MgCl2. DNA was extracted using Phenol/Chloroform/Isoamyl alcohol and precipitated. Tagmented DNA libraries were generated and sequenced using the Illumina NovaSeq 6000 platform.

### CUT&Tag processing

Raw sequencing files were first trimmed of adaptors using fastp^56^ and aligned to the hg38 human genome assembly using bowtie2. Duplicates, non-uniquely aligned reads, and Q scores < 10 were filtered and only proper pairs were retained for analysis. Peaks were called using SEACR^57^ using the -relaxed option and normalized using IgG controls. BedGraph and BigWig files were generated using deepTools and BEDtools^58,59^. Super-enhancers were computed using the ROSE software using H3K27ac BAM files as input^36^. Motif analysis was performed using HOMER for the central 1kb of peaks and compared to known motifs^60^.

### RNA-seq

Raw RNA-seq files were used from data previously generated^12,61^, trimmed of adaptors using fastp, and mapped to the human genome assembly (hg38) using STAR^62^. DEGs were identified using DESeq2^63^. Data was log2 or z-score normalized and noted in the text.

### Luciferase assay

Three candidate regions were selected for enhancer validation using a luciferase reporter assay. Regions were identified by intersecting pcHi-C chromatin loops with H3K27ac peaks obtained from CUT&Tag experiments. Each ∼1 kb region was synthesized as a gBlocks Gene Fragment (Integrated DNA Technologies, IDT) and cloned into the pGL4.23 [luc2/minP] vector (Promega, E841A), which contains the firefly luciferase reporter gene. Constructs were co-transfected with a renilla luciferase control plasmid (E691A) at a 1:10 molar ratio into fetal skeletal muscle progenitor cells (SMPCs). Successful cloning was confirmed by Sanger sequencing. Luciferase activity was measured using the Dual-Luciferase Reporter Assay System (Promega, E1910) on a VICTOR Nivo Multimode Microplate Reader. Firefly luciferase signals were normalized to renilla luciferase, and relative enhancer activity was expressed as fold change compared with a control DNA region (scramble, chr9:6,161,550–6,162,024) lacking active chromatin marks. For each assay, cells were transfected with each construct in triplicate, and three independent experiments were performed. To identify significant increases in luciferase activity, the mean luciferase signal for each construct was compared to that of the control within each technical replicate (n = 3) using a one-tailed paired t-test, with significance defined as P < 0.05.

### CRISPRa experiments and qRT-PCR

sgRNAs for CRISPRa experiments were designed using IDT’s CRISPR Design Tools. Tiled guides targeting respective enhancers (1, 2, or 3) were transfected along with CRISPR/dCas9-VPR mRNA into primary fetal-SMPCs using lipofectamine (MessengerMax). Primary fetal-SMPCs were isolated via FACS as described above and plated in 96 well plates. Cells were plated for 24hrs to allow for recovery after FACS prior to treatment. Cells were treated for 48hrs and then collected for qPCR analysis for *PAX7*. Total RNA was extracted (Qiagen RNease Micro kit) according to manufacter’s instructions. cDNA was synthesized using the iScript Reverse Transcription Supermix and qRT-PCR was performed using SsoAdvanced Universal SYBR Green Supermix with technical triplicates on a Thermo Fisher Scientific QuantStudio 6 Pro Real-Time PCR System. All primer pairs were selected from PrimerBank for *PAX7* and *RPL13A*, as previously described^64,65^.

## Acknowledgements

We thank current and former members of the Pyle lab (M. Hicks and H. Xi) for their help with reagent preparation and flow cytometry feedback. We would also like to thank the Flow Cytometry and High Throughput Sequencing cores from the Eli and Edythe Broad Center of Regenerative Medicine and Stem Cell Research (BSCRC) at the University of California, Los Angeles (UCLA). We acknowledge the Technology Center for Genomics & Bioinformatics, Translational Pathology, and Janis V. Giorgi Flow Cytometry cores at the UCLA Jonsson Comprehensive Cancer Center (JCCC). A.D.P. is funded by CIRM Quest (DISC2-10696), the National Institute of Arthritis and Musculoskeletal and Skin Diseases (NIAMS) (R01 AR064327), UCLA BSCRC, the Rose Hills Foundation, the Ablon Scholars Award and the George and Nouhad Ayoub Centennial Chair for Life Science Innovation. M.A.R. is funded by the University of California President’s Postdoctoral Fellowship Program and the Ford Foundation Postdoctoral Fellowship program. D.R.S. is funded by ADA (1-24-ACE-21), and NIDDK grant U24DK132746-01, UCLA LIFT-UP and National Institutes of Health Office of Disease Prevention (ODP).

## Author contributions

M.A.R., and A.D.P. conceived and oversaw the project. M.A.R., P.C., H.L.L, G.C., E.S., K.S., D.G., L.G., D-H.H., L.C. performed the experiments, M.A.R., C.N., H.L.L, D.R.S. analyzed the data. M.A.R. and A.D.P. wrote the manuscript with editing from C.N., D.R.S., P.L.P. with additional edits and approval from all authors.

## Supplementary figures and figure legends

**Figure S1.**
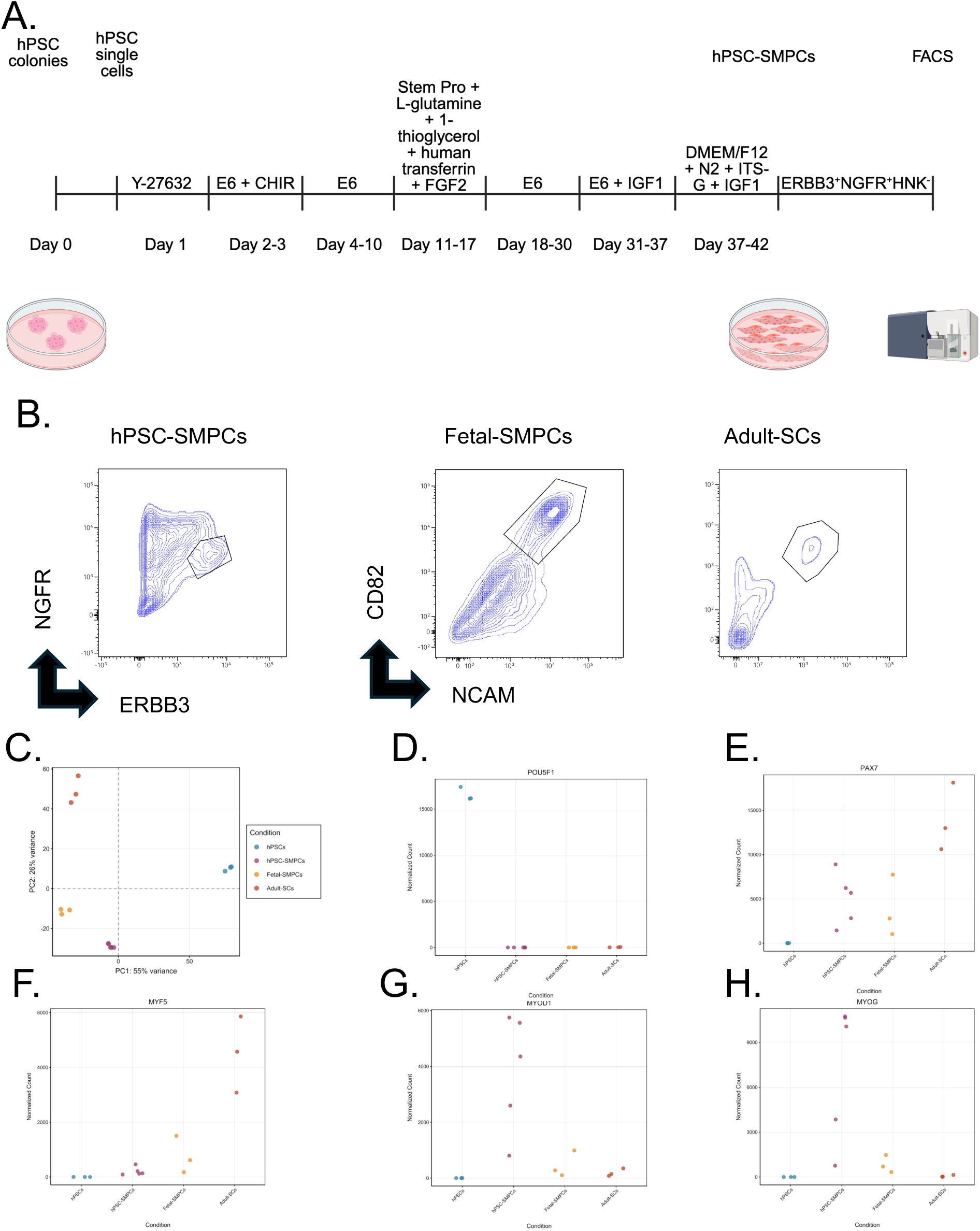
Characterization of myogenic progenitors and adult-SCs. **A)** Differentiation timeline of *in-vitro* derived hPSC-SMPCs. **B)** FACS plots of myogenic progenitors and adult-SCs. **C)** PCA plot depicting transcriptional differences across our samples. **D-H)** Normalized transcriptional plot counts for known transcriptional regulators of across hPSCs and myogenic lineages.

**Figure S2.**
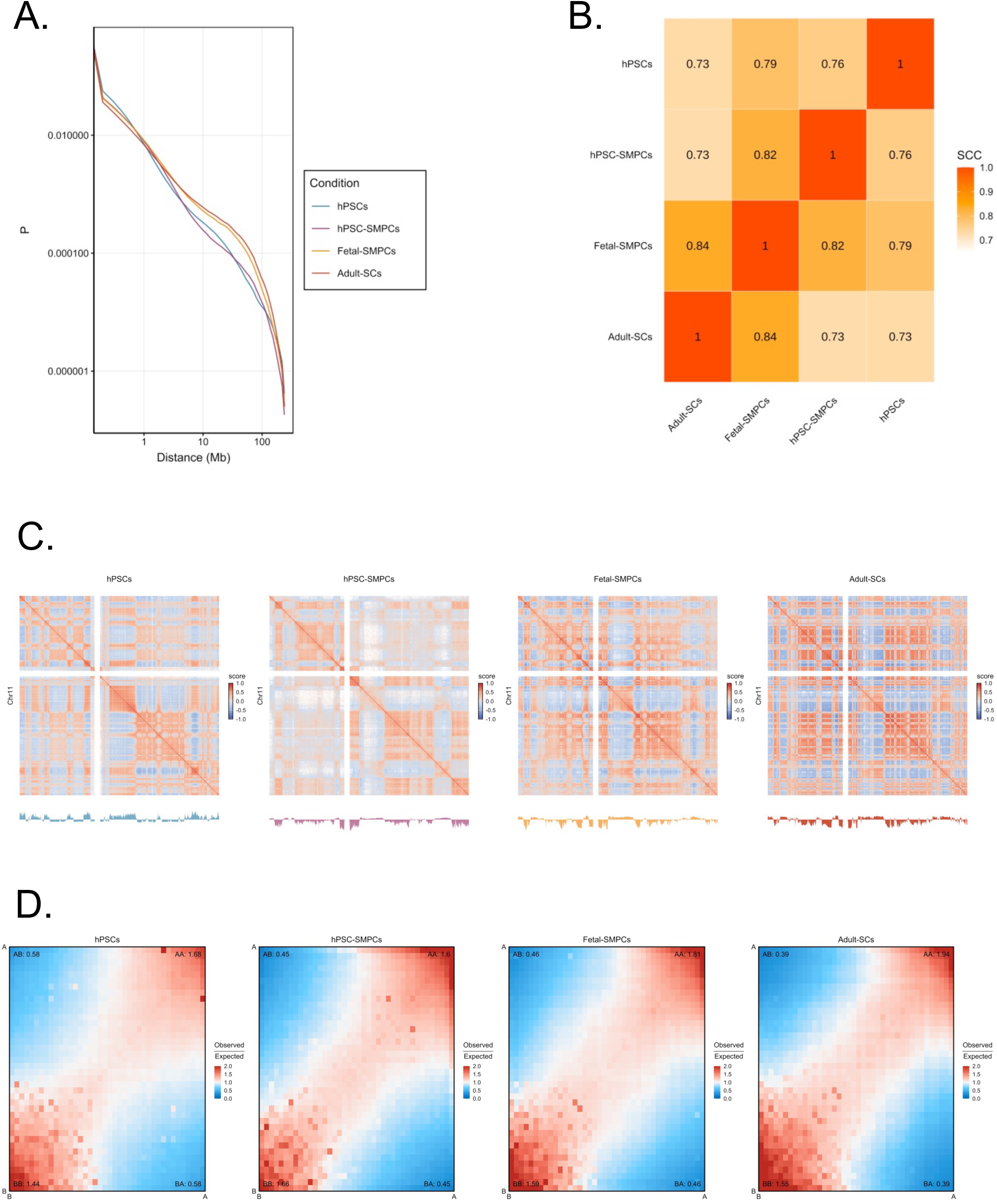
Comparison of long-range interactions and compartmentalization. **A)** Interaction probability across increasing distance in hPSCs, hPSC-SMPCs, fetal-SMPCs, and adult-SCs. **B)** Correlation of Hi-C across samples using HiCRep^48^. **C)** Compartmentalization of chromosome 11 across conditions (top matrix) with associated principal component 1 values calculated using dcHiC^49^. **D)** Genome-wide saddle plots depicting compartment strengths across all four conditions. Data was visualized using GENOVA^50^.

**Figure S3.**
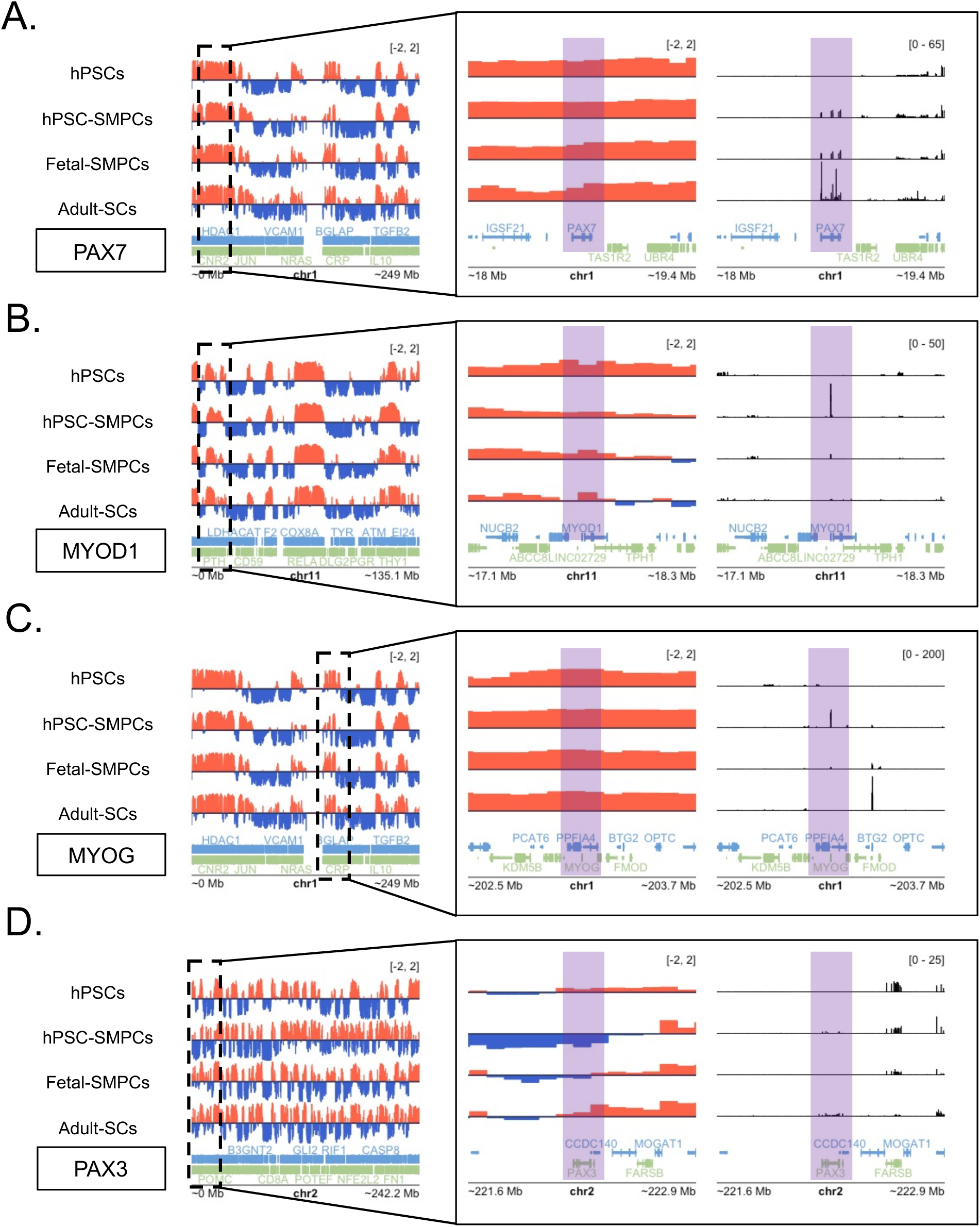
Compartmentalization of myogenic transcription factors across conditions. A-D) Compartmentalization of transcription factors that control satellite cell specification and/or differentiation across all four conditions. Surrounding chromatin for each respective gene is first visualize and subsequently zoomed in to visualize the immediate surrounding regions. Positive red values depict the A compartment, while negative blue values depict B compartments. All compartment data was analyzed using dcHiC^49^ and visualized using Plotgardener^51^.

**Figure S4.**
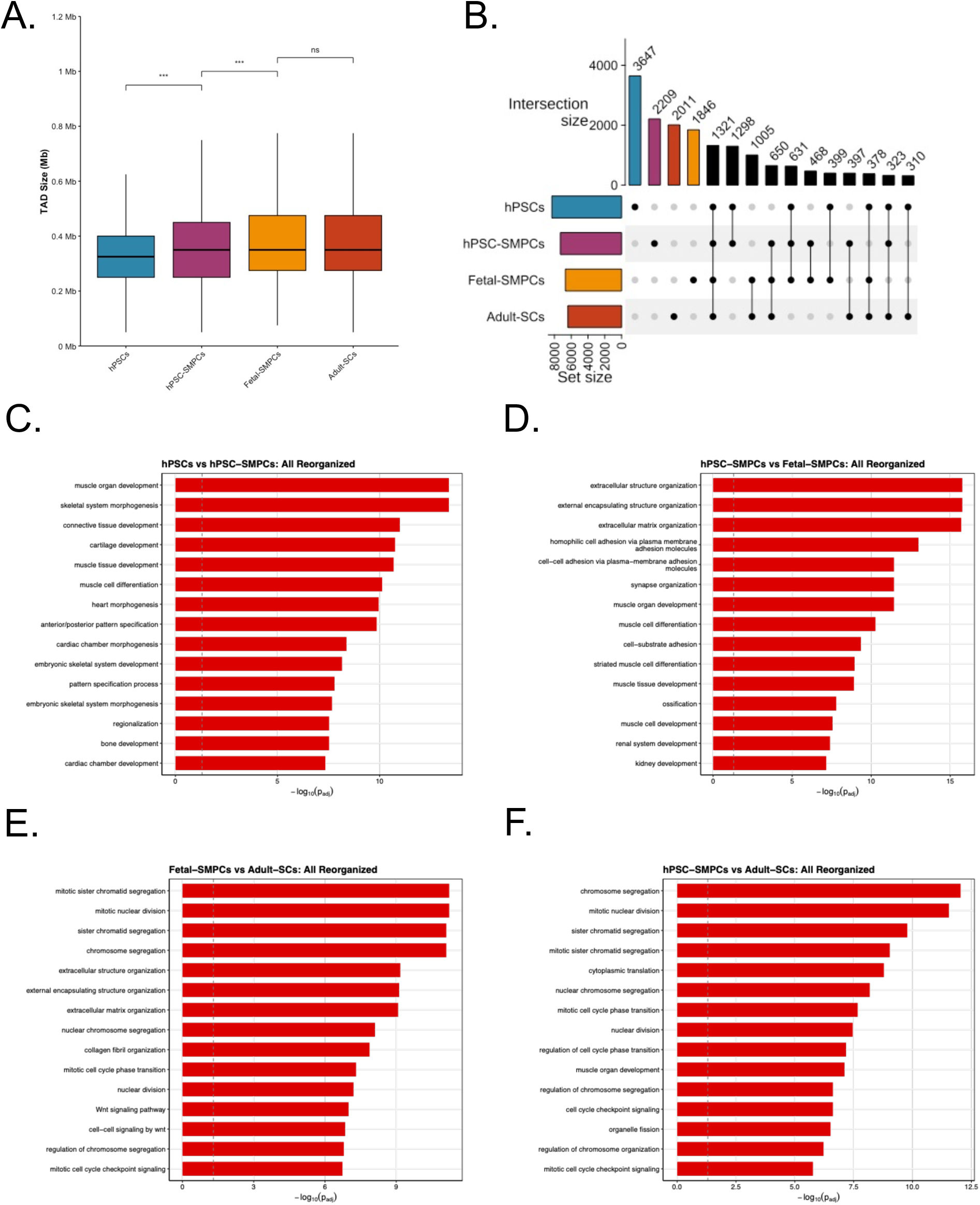
TAD characterization and reorganization across conditions. **A)** TAD size across all conditions. **B)** Quantifying unique TADs and conservation of TADs across conditions. **C-F)** Pairwise GO Analysis on DEGs within all reorganized TADs. Mann-Whitney U test was used for significance testing. Significance is displayed as *p < 0.05, **p < 0.001, ***p < 0.0001, ****p < 0.00001.

**Figure S5.**
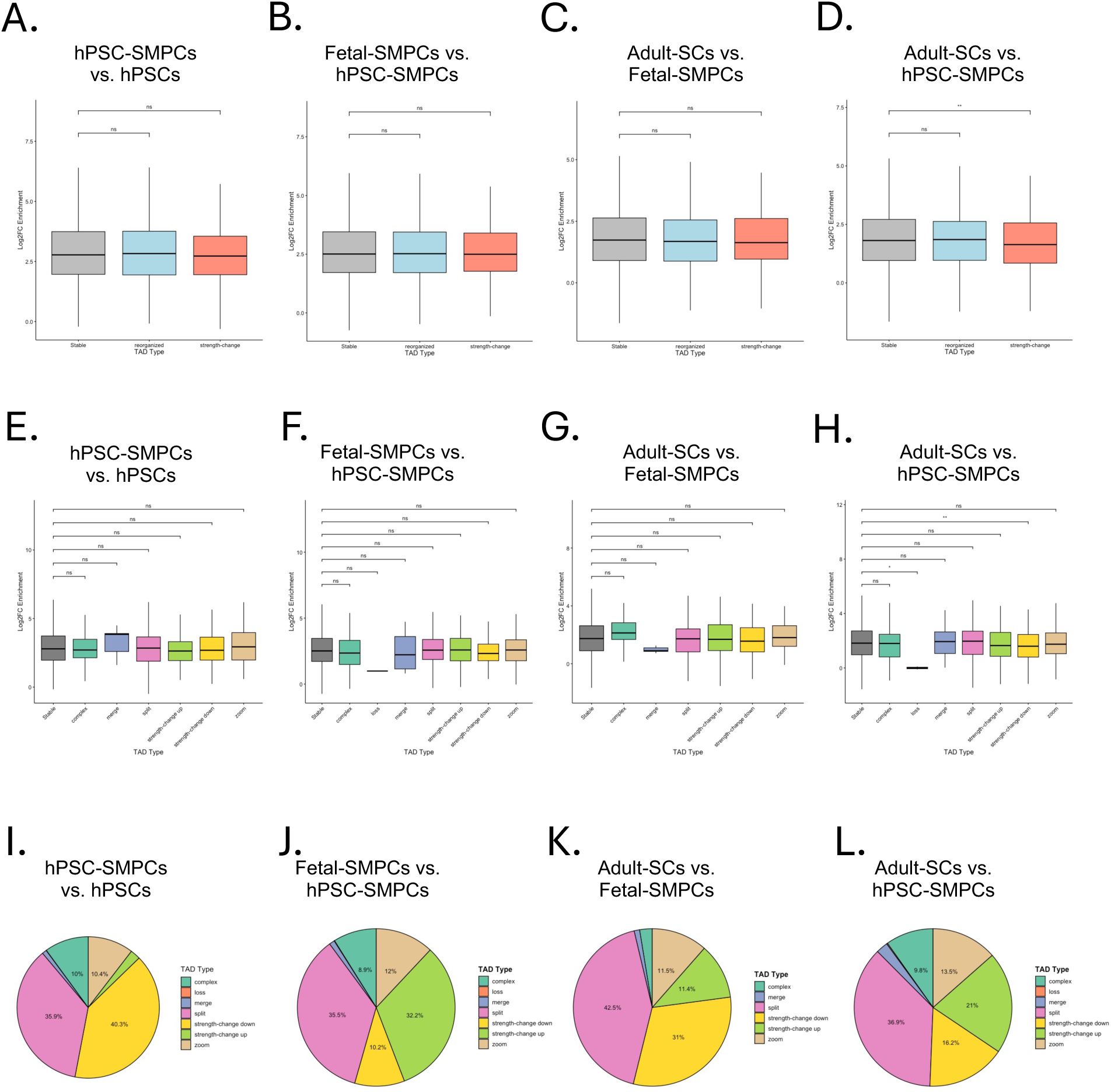
CTCF enrichment at reorganized TADs and genes within reorganized TADs. A-D) Comparing CTCF enrichment at Collapsed TAD reorganization subtypes at later stage vs. previous stage. Reorganization subtypes were grouped together as stable, strength-change (up or down), or otherwise reorganized. **E-H)** Comparing CTCF enrichment at specific reorganization subtypes at later stage vs. previous stage. **I-L)** Comparing proportion of genes that were reorganized within TADs at later stage vs. earlier stage. Mann-Whitney U test was used for significance testing. Significance is displayed as *p < 0.05, **p < 0.001.

**Figure S6.**
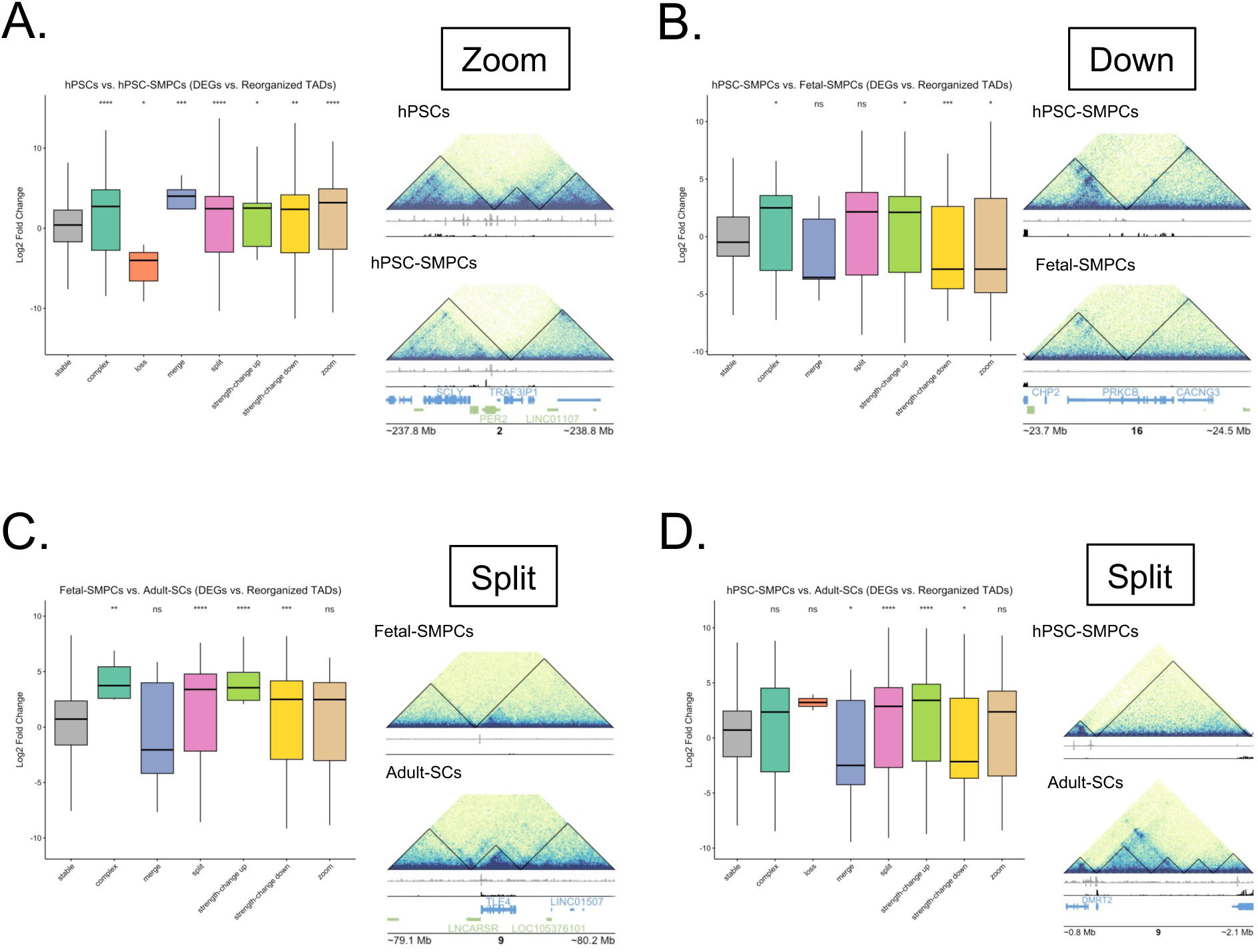
DEGs at Reorganized TAD subtypes. **A)** Depicting gene expression changes between hPSCs and hPSC-SMPCs across reorganization subtypes with an example of a zoom reorganization event around the *PER2* gene showing higher expression of *PER2* in hPSC-SMPCs in the reorganized TAD compared to hPSCs. **B)** Gene expression changes across reorganized TAD subtypes between hPSC-SMPCs and fetal-SMPCs around the *PRKCB* gene, which is decreased in fetal-SMPCs compared to hPSC-SMPCs. **C)** Gene expression changes withing reorganized TAD subtypes between fetal-SMPCs and adult-SCs with an example of a split TAD around *TLE4* showing increased expression in adult-SCs. **D)** Gene expression changes withing reorganized TAD subtypes between hPSC-SMPCs and adult-SCs. An example of a split TAD subtype is depicted around *DMRT2*, which shows higher expression in adult-SCs as compared to hPSC-SMPCs. Mann-Whitney U test was used for significance testing. Significance is displayed as *p < 0.05, **p < 0.001, ***p < 0.0001, ****p < 0.00001.

**Figure S7.**
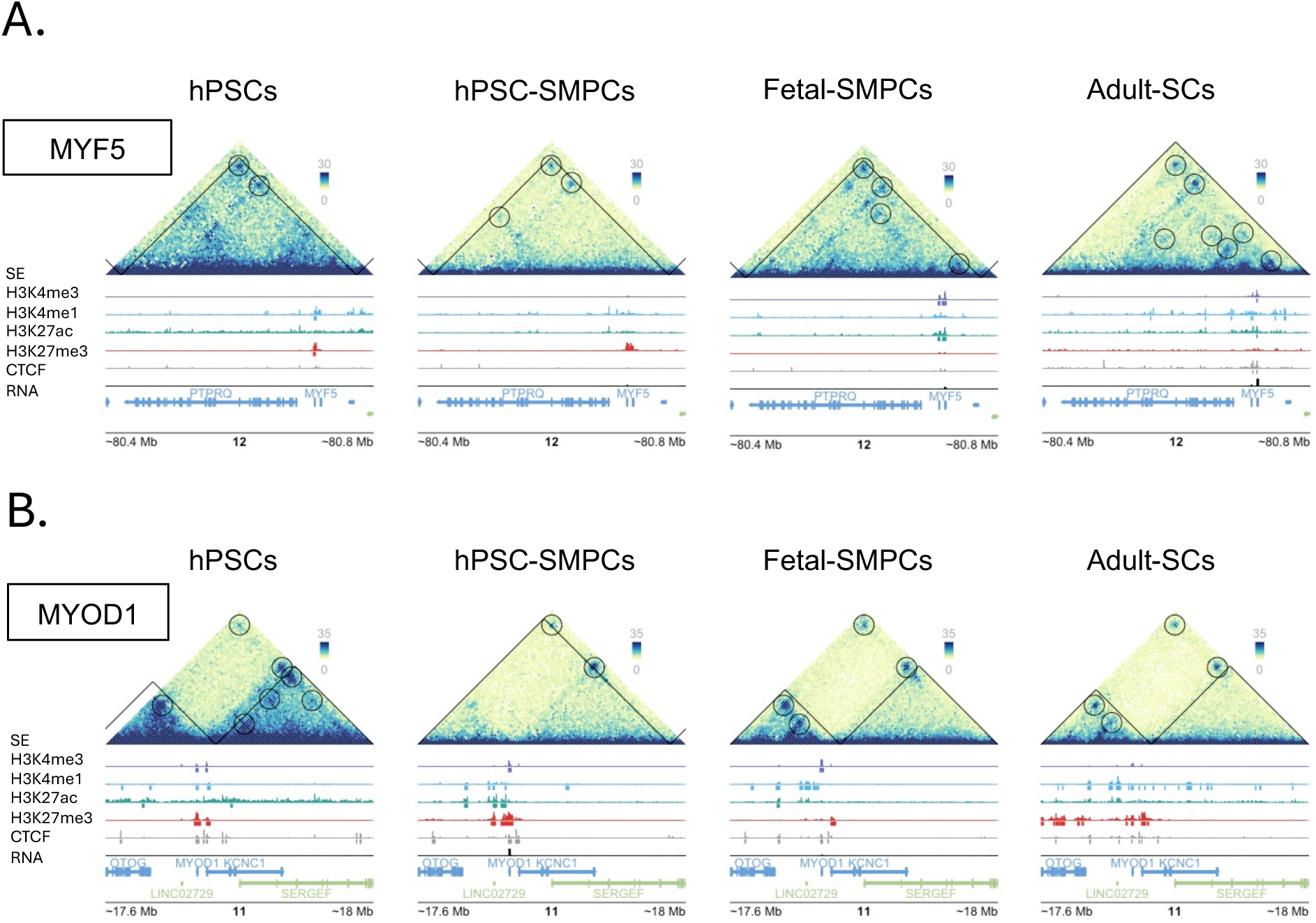
Chromatin architecture and looping at *MYF5* and *MYOD1* across development. TAD organization and chromatin loop formation at **A)** *MYF5* and **B)** *MYOD1*. TADs are conserved across development for *MYF5*, while hPSC-SMPCs shows TAD reorganization followed by maintenance in fetal-SMPCs and adult-SCs for *MYOD1*. Chromatin loops show dynamic behavior for *MYF5* and *MYOD1* with chromatin loops having an additive effect on *MYF5* expression toward adult-SCs, while *MYOD1* expression is only seen in hPSC-SMPCs when chromatin loops have the lowest count. Genome tracks for CUT&Tag and RNA-seq on bottom of pyramid plots.

**Figure S8.**
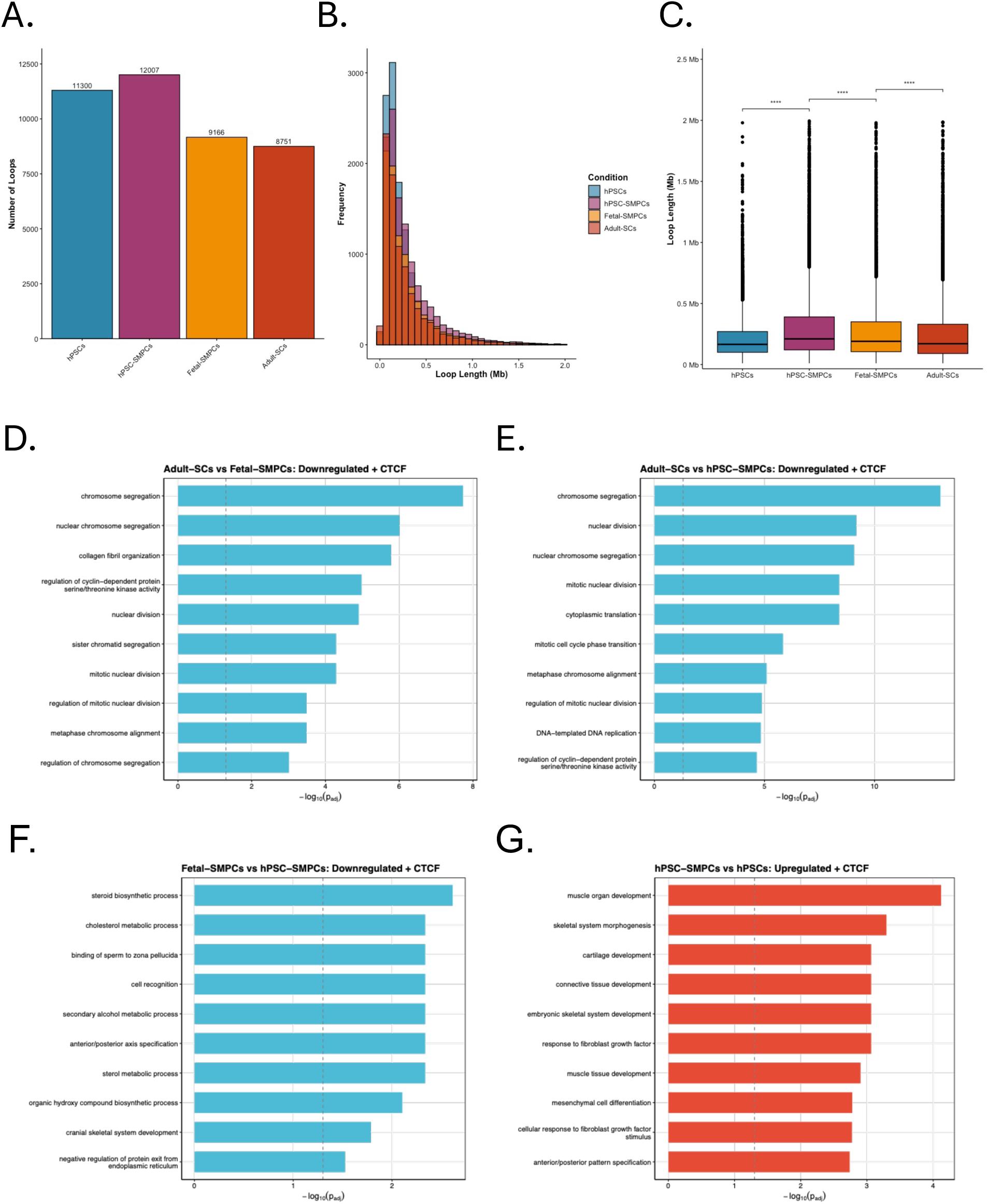
Chromatin loops and CTCF enrichment across development. **A)** Number of chromatin loops called using HiCExplorer^27^. **B)** Histogram of chromatin loop sizes across all four conditions and quantified in **C)**. **D-G)** Pairwise GO Analysis on down- or upregulated DEGs with CTCF enriched at the promoter.

**Figure S9.**
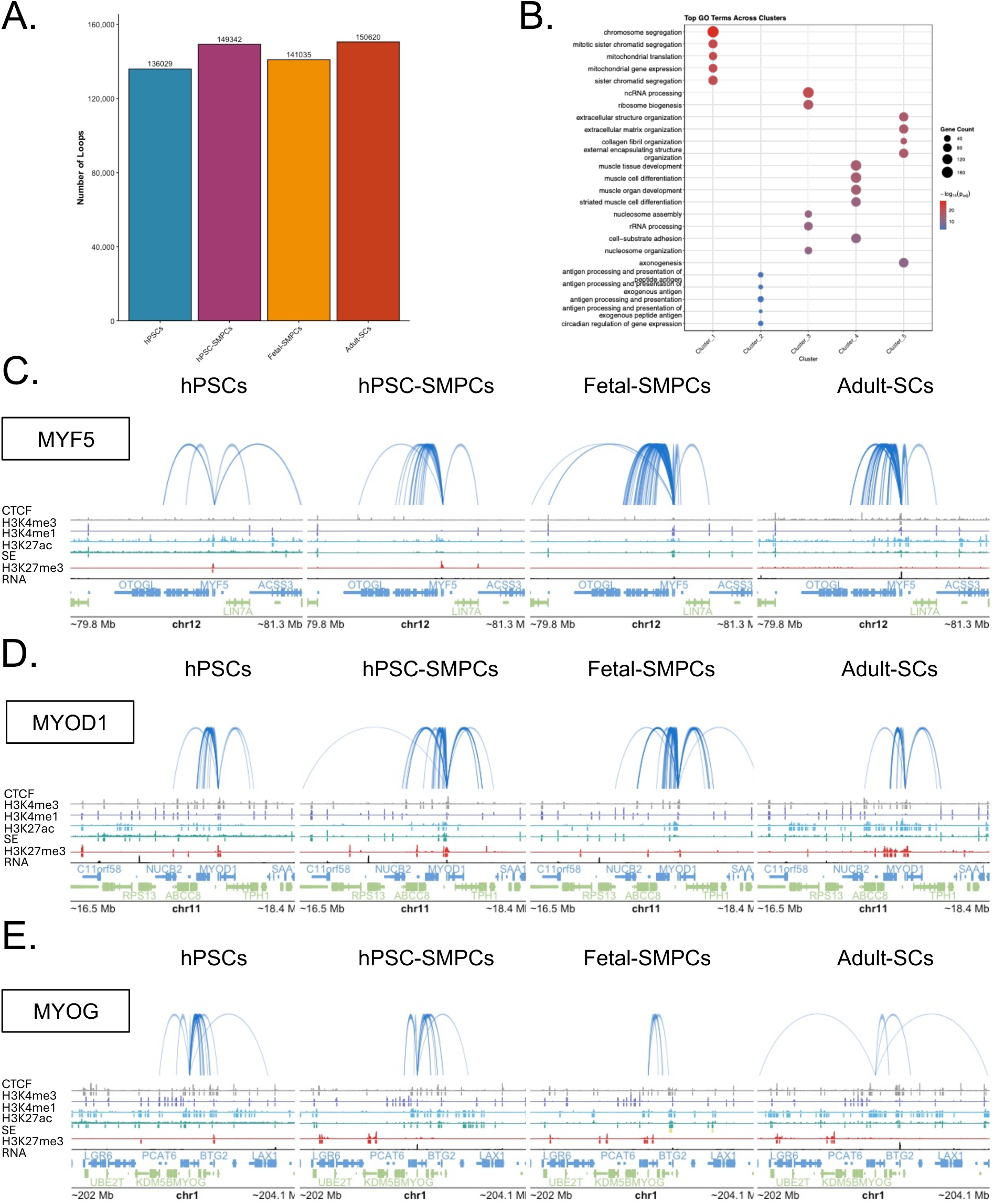
Promoter-capture Hi-C and enhancer integration across myogenic relevant transcription factors. **A)** Total number of loops called from pcHi-C experiments analyzed using CHiAGO^52^. **B)** GO analysis on DEGs intersected with H3K27ac anchored pcHiC loops. Each cluster depicts top 5 enriched GO terms. **C-E)** Arc plot at *MYF5*, *MYOD1*, *MYOG*, loci depicting pcHi-C interactions across all four conditions along with genome tracks from CUT&Tag and RNA-seq data.

## References

1. Sacco, A., Doyonnas, R., Kraft, P., Vitorovic, S., and Blau, H. Self-renewal and expansion of single transplanted muscle stem cells. 456. 10.1038/nature07384.

2. Seale, P., Sabourin, L.A., Girgis-Gabardo, A., Mansouri, A., Gruss, P., and Rudnicki, M.A. (2000). Pax7 Is Required for the Specification of Myogenic Satellite Cells. Cell 102, 777–786. 10.1016/s0092-8674(00)00066-0.

3. Furlong, E.E., and Levine, M. (2018). Developmental enhancers and chromosome topology. Science 361, 1341–1345. 10.1126/science.aau0320.

4. Pollex, T., Rabinowitz, A., Gambetta, M.C., Marco-Ferreres, R., Viales, R.R., Jankowski, A., Schaub, C., and Furlong, E.E.M. (2024). Enhancer–promoter interactions become more instructive in the transition from cell-fate specification to tissue differentiation. Nat. Genet., 1–11. 10.1038/s41588-024-01678-x.

5. Lieberman-Aiden, E., Berkum, N.L. van, Williams, L., Imakaev, M., Ragoczy, T., Telling, A., Amit, I., Lajoie, B.R., Sabo, P.J., Dorschner, M.O., et al. (2009). Comprehensive Mapping of Long-Range Interactions Reveals Folding Principles of the Human Genome. Science 326, 289–293. 10.1126/science.1181369.

6. Rao, S.S.P., Huntley, M.H., Durand, N.C., Stamenova, E.K., Bochkov, I.D., Robinson, J.T., Sanborn, A.L., Machol, I., Omer, A.D., Lander, E.S., et al. (2014). A 3D Map of the Human Genome at Kilobase Resolution Reveals Principles of Chromatin Looping. Cell 159, 1665–1680. 10.1016/j.cell.2014.11.021.

7. Dixon, J.R., Selvaraj, S., Yue, F., Kim, A., Li, Y., Shen, Y., Hu, M., Liu, J.S., and Ren, B. (2012). Topological domains in mammalian genomes identified by analysis of chromatin interactions. Nature 485, 376. 10.1038/nature11082.

8. Zhang, N., Mendieta-Esteban, J., Magli, A., Lilja, K.C., Perlingeiro, R.C.R., Marti-Renom, M.A., Tsirigos, A., and Dynlacht, B.D. (2020). Muscle progenitor specification and myogenic differentiation are associated with changes in chromatin topology. Nat Commun 11, 6222. 10.1038/s41467-020-19999-w.

9. Zhao, Y., Ding, Y., He, L., Zhou, Q., Chen, X., Li, Y., Alfonsi, M.V., Wu, Z., Sun, H., and Wang, H. (2023). Multiscale 3D genome reorganization during skeletal muscle stem cell lineage progression and aging. Sci. Adv. 9, eabo1360. 10.1126/sciadv.abo1360.

10. Khateb, M., Perovanovic, J., Ko, K.D., Jiang, K., Feng, X., Acevedo-Luna, N., Chal, J., Ciuffoli, V., Genzor, P., Simone, J., et al. (2022). Transcriptomics, regulatory syntax, and enhancer identification in mesoderm-induced ESCs at single-cell resolution. Cell Reports 40, 111219. 10.1016/j.celrep.2022.111219.

11. Xi, H., Langerman, J., Sabri, S., Chien, P., Young, C.S., Younesi, S., Hicks, M., Gonzalez, K., Fujiwara, W., Marzi, J., et al. (2020). A Human Skeletal Muscle Atlas Identifies the Trajectories of Stem and Progenitor Cells across Development and from Human Pluripotent Stem Cells. Cell Stem Cell. 10.1016/j.stem.2020.04.017.

12. Hicks, M.R., Hiserodt, J., Paras, K., Fujiwara, W., Eskin, A., Jan, M., Xi, H., Young, C.S., Evseenko, D., Nelson, S.F., et al. (2018). ERBB3 and NGFR mark a distinct skeletal muscle progenitor cell in human development and hPSCs. Nature Cell Biology 20, 46–57. 10.1038/s41556-017-0010-2.

13. Shelton, M., Metz, J., Liu, J., Carpenedo, R.L., Demers, S.-P., Stanford, W.L., and Skerjanc, I.S. (2014). Derivation and Expansion of PAX7-Positive Muscle Progenitors from Human and Mouse Embryonic Stem Cells. Stem Cell Rep 3, 516–529. 10.1016/j.stemcr.2014.07.001.

14. Alonso-Martin, S., Rochat, A., Mademtzoglou, D., Morais, J., Reyniès, A. de, Auradé, F., Chang, T.H.-T., Zammit, P.S., and Relaix, F. (2016). Gene Expression Profiling of Muscle Stem Cells Identifies Novel Regulators of Postnatal Myogenesis. Front. Cell Dev. Biol. 4, 58. 10.3389/fcell.2016.00058.

15. Dixon, J.R., Jung, I., Selvaraj, S., Shen, Y., Antosiewicz-Bourget, J.E., Lee, A., Ye, Z., Kim, A., Rajagopal, N., Xie, W., et al. (2015). Chromatin architecture reorganization during stem cell differentiation. Nature 518, 331. 10.1038/nature14222.

16. Takayama, N., Murison, A., Takayanagi, S., Arlidge, C., Zhou, S., Garcia-Prat, L., Chan-Seng-Yue, M., Zandi, S., Gan, O.I., Boutzen, H., et al. (2020). The Transition from Quiescent to Activated States in Human Hematopoietic Stem Cells Is Governed by Dynamic 3D Genome Reorganization. Cell Stem Cell. 10.1016/j.stem.2020.11.001.

17. Stik, G., Vidal, E., Barrero, M., Cuartero, S., Vila-Casadesús, M., Mendieta-Esteban, J., Tian, T.V., Choi, J., Berenguer, C., Abad, A., et al. (2020). CTCF is dispensable for immune cell transdifferentiation but facilitates an acute inflammatory response. Nat Genet, 1–7. 10.1038/s41588-020-0643-0.

18. Nora, E.P., Goloborodko, A., Valton, A.-L., Gibcus, J.H., Uebersohn, A., Abdennur, N., Dekker, J., Mirny, L.A., and Bruneau, B.G. (2017). Targeted Degradation of CTCF Decouples Local Insulation of Chromosome Domains from Genomic Compartmentalization. Cell 169, 930–944.e22. 10.1016/j.cell.2017.05.004.

19. Li, Y., Nakka, K., Olender, T., Gingras-Gelinas, P., Wong, M.M.-K., Robinson, D.C.L., Bandukwala, H., Palii, C.G., Neyret, O., Brand, M., et al. (2021). Chromatin and transcription factor profiling in rare stem cell populations using CUT&Tag. Star Protoc 2, 100751. 10.1016/j.xpro.2021.100751.

20. Robinson, D.C.L., Ritso, M., Nelson, G.M., Mokhtari, Z., Nakka, K., Bandukwala, H., Goldman, S.R., Park, P.J., Mounier, R., Chazaud, B., et al. (2021). Negative elongation factor regulates muscle progenitor expansion for efficient myofiber repair and stem cell pool repopulation. Dev Cell 56, 1014–1029.e7. 10.1016/j.devcel.2021.02.025.

21. Kaya-Okur, H.S., Wu, S.J., Codomo, C.A., Pledger, E.S., Bryson, T.D., Henikoff, J.G., Ahmad, K., and Henikoff, S. (2019). CUT&Tag for efficient epigenomic profiling of small samples and single cells. Nat Commun 10, 1930. 10.1038/s41467-019-09982-5.

22. Chang, L., and Noordermeer, D. (2024). Permeable TAD boundaries and their impact on genome-associated functions. BioEssays 46, e2400137. 10.1002/bies.202400137.

23. Chang, L.-H., Ghosh, S., and Noordermeer, D. (2020). TADs and Their Borders: Free Movement or Building a Wall? J Mol Biol 432, 643–652. 10.1016/j.jmb.2019.11.025.

24. Narendra, V., Bulajić, M., Dekker, J., Mazzoni, E.O., and Reinberg, D. (2016). CTCF-mediated topological boundaries during development foster appropriate gene regulation. Genes Dev. 30, 2657–2662. 10.1101/gad.288324.116.

25. Hua, D., Gu, M., Zhang, X., Du, Y., Xie, H., Qi, L., Du, X., Bai, Z., Zhu, X., and Tian, D. (2024). DiffDomain enables identification of structurally reorganized topologically associating domains. Nat. Commun. 15, 502. 10.1038/s41467-024-44782-6.

26. Yoshioka, K., Nagahisa, H., Miura, F., Araki, H., Kamei, Y., Kitajima, Y., Seko, D., Nogami, J., Tsuchiya, Y., Okazaki, N., et al. (2021). Hoxa10 mediates positional memory to govern stem cell function in adult skeletal muscle. Sci Adv 7, eabd7924. 10.1126/sciadv.abd7924.

27. Wolff, J., Rabbani, L., Gilsbach, R., Richard, G., Manke, T., Backofen, R., and Grüning, B.A. (2020). Galaxy HiCExplorer 3: a web server for reproducible Hi-C, capture Hi-C and single-cell Hi-C data analysis, quality control and visualization. Nucleic Acids Res. 10.1093/nar/gkaa220.

28. Song, Y., Liang, Z., Zhang, J., Hu, G., Wang, J., Li, Y., Guo, R., Dong, X., Babarinde, I.A., Ping, W., et al. (2022). CTCF functions as an insulator for somatic genes and a chromatin remodeler for pluripotency genes during reprogramming. Cell Reports 39, 110626. 10.1016/j.celrep.2022.110626.

29. Murphy, D., Salataj, E., Giammartino, D.C.D., Rodriguez-Hernaez, J., Kloetgen, A., Garg, V., Char, E., Uyehara, C.M., Ee, L., Lee, U., et al. (2023). 3D Enhancer–promoter networks provide predictive features for gene expression and coregulation in early embryonic lineages. Nat. Struct. Mol. Biol., 1–16. 10.1038/s41594-023-01130-4.

30. Wang, R., Chen, F., Chen, Q., Wan, X., Shi, M., Chen, A.K., Ma, Z., Li, G., Wang, M., Ying, Y., et al. (2022). MyoD is a 3D genome structure organizer for muscle cell identity. Nat Commun 13, 205. 10.1038/s41467-021-27865-6.

31. Dall’Agnese, A., Caputo, L., Nicoletti, C., Iulio, J. di, Schmitt, A., Gatto, S., Diao, Y., Ye, Z., Forcato, M., Perera, R., et al. (2019). Transcription Factor-Directed Re-wiring of Chromatin Architecture for Somatic Cell Nuclear Reprogramming toward trans-Differentiation. Mol Cell 76. 10.1016/j.molcel.2019.07.036.

32. Maltzahn, J. von, Jones, A.E., Parks, R.J., and Rudnicki, M.A. (2013). Pax7 is critical for the normal function of satellite cells in adult skeletal muscle. Proc National Acad Sci 110, 16474–16479. 10.1073/pnas.1307680110.

33. Hansen, A.S., Pustova, I., Cattoglio, C., Tjian, R., and Darzacq, X. (2017). CTCF and cohesin regulate chromatin loop stability with distinct dynamics. Elife 6, e25776. 10.7554/elife.25776.

34. Yoon, S., Chandra, A., and Vahedi, G. (2022). Stripenn detects architectural stripes from chromatin conformation data using computer vision. Nat. Commun. 13, 1602. 10.1038/s41467-022-29258-9.

35. Vian, L., Pękowska, A., Rao, S., Kieffer-Kwon, K.-R., Jung, S., Baranello, L., Huang, S.-C., Khattabi, L., Dose, M., Pruett, N., et al. (2018). The Energetics and Physiological Impact of Cohesin Extrusion. Cell 173, 1165–1178.e20. 10.1016/j.cell.2018.03.072.

36. Whyte, W.A., Orlando, D.A., Hnisz, D., Abraham, B.J., Lin, C.Y., Kagey, M.H., Rahl, P.B., Lee, T., and Young, R.A. (2013). Master Transcription Factors and Mediator Establish Super-Enhancers at Key Cell Identity Genes. Cell 153, 307–319. 10.1016/j.cell.2013.03.035.

37. Mifsud, B., Tavares-Cadete, F., Young, A.N., Sugar, R., Schoenfelder, S., Ferreira, L., Wingett, S.W., Andrews, S., Grey, W., Ewels, P.A., et al. (2015). Mapping long-range promoter contacts in human cells with high-resolution capture Hi-C. Nat Genet 47, ng.3286. 10.1038/ng.3286.

38. Sobreira, D.R., Joslin, A.C., Zhang, Q., Williamson, I., Hansen, G.T., Farris, K.M., Sakabe, N.J., Sinnott-Armstrong, N., Bozek, G., Jensen-Cody, S.O., et al. (2021). Extensive pleiotropism and allelic heterogeneity mediate metabolic effects of IRX3 and IRX5. Science 372, 1085–1091. 10.1126/science.abf1008.

39. Dong, A., Liu, J., Lin, K., Zeng, W., So, W.-K., Hu, S., and Cheung, T.H. (2022). Global chromatin accessibility profiling analysis reveals a chronic activation state in aged muscle stem cells. Iscience 25, 104954. 10.1016/j.isci.2022.104954.

40. Villar, D., Berthelot, C., Aldridge, S., Rayner, T.F., Lukk, M., Pignatelli, M., Park, T.J., Deaville, R., Erichsen, J.T., Jasinska, A.J., et al. (2015). Enhancer Evolution across 20 Mammalian Species. Cell 160, 554–566. 10.1016/j.cell.2015.01.006.

41. Schmidt, D., Schwalie, P.C., Wilson, M.D., Ballester, B., Gonçalves, Â., Kutter, C., Brown, G.D., Marshall, A., Flicek, P., and Odom, D.T. (2012). Waves of Retrotransposon Expansion Remodel Genome Organization and CTCF Binding in Multiple Mammalian Lineages. Cell 148, 335–348. 10.1016/j.cell.2011.11.058.

42. Machado, L., Lima, J. de, Fabre, O., Proux, C., Legendre, R., Szegedi, A., Varet, H., Ingerslev, L., Barrès, R., Relaix, F., et al. (2017). In Situ Fixation Redefines Quiescence and Early Activation of Skeletal Muscle Stem Cells. Cell Reports 21, 1982–1993. 10.1016/j.celrep.2017.10.080.

43. Chang, L.-H., Ghosh, S., Papale, A., Luppino, J.M., Miranda, M., Piras, V., Degrouard, J., Edouard, J., Poncelet, M., Lecouvreur, N., et al. (2023). Multi-feature clustering of CTCF binding creates robustness for loop extrusion blocking and Topologically Associating Domain boundaries. Nat. Commun. 14, 5615. 10.1038/s41467-023-41265-y.

44. Long, H.S., Greenaway, S., Powell, G., Mallon, A.-M., Lindgren, C.M., and Simon, M.M. (2022). Making sense of the linear genome, gene function and TADs. Epigenet Chromatin 15, 4. 10.1186/s13072-022-00436-9.

45. Wu, H.-J., Landshammer, A., Stamenova, E.K., Bolondi, A., Kretzmer, H., Meissner, A., and Michor, F. (2021). Topological isolation of developmental regulators in mammalian genomes. Nat Commun 12, 4897. 10.1038/s41467-021-24951-7.

46. Gryder, B.E., Wachtel, M., Chang, K., Demerdash, O.E., Aboreden, N.G., Mohammed, W., Ewert, W., Pomella, S., Rota, R., Wei, J.S., et al. (2020). Miswired enhancer logic drives a cancer of the muscle lineage. Iscience 23, 101103. 10.1016/j.isci.2020.101103.

47. Gryder, B.E., Pomella, S., Sayers, C., Wu, X.S., Song, Y., Chiarella, A.M., Bagchi, S., Chou, H.-C., Sinniah, R.S., Walton, A., et al. (2019). Histone hyperacetylation disrupts core gene regulatory architecture in rhabdomyosarcoma. Nat Genet 51, 1714–1722. 10.1038/s41588-019-0534-4.

48. Yang, T., Zhang, F., Yardımcı, G.G., Song, F., Hardison, R.C., Noble, W.S., Yue, F., and Li, Q. (2017). HiCRep: assessing the reproducibility of Hi-C data using a stratum-adjusted correlation coefficient. Genome Res 27, 1939–1949. 10.1101/gr.220640.117.

49. Chakraborty, A., Wang, J.G., and Ay, F. (2022). dcHiC detects differential compartments across multiple Hi-C datasets. Nat Commun 13, 6827. 10.1038/s41467-022-34626-6.

50. 50. van der Weide, R.H., van den Brand, T., Haarhuis, J.H.I., Teunissen, H., Rowland, B.D., and de Wit, E. (2021). Hi-C analyses with GENOVA: a case study with cohesin variants. Nar Genom Bioinform 3, lqab040-. 10.1093/nargab/lqab040.

51. Kramer, N.E., Davis, E.S., Wenger, C.D., Deoudes, E.M., Parker, S.M., Love, M.I., and Phanstiel, D.H. (2022). Plotgardener: cultivating precise multi-panel figures in R. Bioinformatics 38, 2042–2045. 10.1093/bioinformatics/btac057.

52. Cairns, J., Freire-Pritchett, P., Wingett, S.W., Várnai, C., Dimond, A., Plagnol, V., Zerbino, D., Schoenfelder, S., Javierre, B.-M., Osborne, C., et al. (2016). CHiCAGO: robust detection of DNA looping interactions in Capture Hi-C data. Genome Biol 17, 127. 10.1186/s13059-016-0992-2.

53. Durand, N.C., Shamim, M.S., Machol, I., Rao, S.S.P., Huntley, M.H., Lander, E.S., and Aiden, E.L. (2016). Juicer Provides a One-Click System for Analyzing Loop-Resolution Hi-C Experiments. Cell Syst 3, 95–98. 10.1016/j.cels.2016.07.002.

54. Flyamer, I.M., Illingworth, R.S., and Bickmore, W.A. (2020). Coolpup.py: versatile pile-up analysis of Hi-C data. Bioinformatics 36, 2980–2985. 10.1093/bioinformatics/btaa073.

55. Wingett, S., Ewels, P., Furlan-Magaril, M., Nagano, T., Schoenfelder, S., Fraser, P., and Andrews, S. (2015). HiCUP: pipeline for mapping and processing Hi-C data. F1000Research 4, 1310. 10.12688/f1000research.7334.1.

56. Chen, S., Zhou, Y., Chen, Y., and Gu, J. (2018). fastp: an ultra-fast all-in-one FASTQ preprocessor. Bioinformatics 34, i884–i890. 10.1093/bioinformatics/bty560.

57. Meers, M.P., Tenenbaum, D., and Henikoff, S. (2019). Peak calling by Sparse Enrichment Analysis for CUT&RUN chromatin profiling. Epigenet Chromatin 12, 42. 10.1186/s13072-019-0287-4.

58. Quinlan, A.R., and Hall, I.M. (2010). BEDTools: a flexible suite of utilities for comparing genomic features. Bioinformatics 26, 841–842. 10.1093/bioinformatics/btq033.

59. Ramírez, F., Dündar, F., Diehl, S., Grüning, B.A., and Manke, T. (2014). deepTools: a flexible platform for exploring deep-sequencing data. Nucleic Acids Res. 42, W187–W191. 10.1093/nar/gku365.

60. Heinz, S., Benner, C., Spann, N., Bertolino, E., Lin, Y.C., Laslo, P., Cheng, J.X., Murre, C., Singh, H., and Glass, C.K. (2010). Simple Combinations of Lineage-Determining Transcription Factors Prime cis-Regulatory Elements Required for Macrophage and B Cell Identities. Mol Cell 38, 576–589. 10.1016/j.molcel.2010.05.004.

61. Gill, K.P., Hung, S.S.C., Sharov, A., Lo, C.Y., Needham, K., Lidgerwood, G.E., Jackson, S., Crombie, D.E., Nayagam, B.A., Cook, A.L., et al. (2016). Enriched retinal ganglion cells derived from human embryonic stem cells. Sci. Rep. 6, 30552. 10.1038/srep30552.

62. Dobin, A., Davis, C.A., Schlesinger, F., Drenkow, J., Zaleski, C., Jha, S., Batut, P., Chaisson, M., and Gingeras, T.R. (2012). STAR: ultrafast universal RNA-seq aligner. Bioinformatics 29, 15–21. 10.1093/bioinformatics/bts635.

63. Love, M.I., Huber, W., and Anders, S. (2014). Moderated estimation of fold change and dispersion for RNA-seq data with DESeq2. Genome Biol. 15, 550. 10.1186/s13059-014-0550-8.

64. Wang, X., Spandidos, A., Wang, H., and Seed, B. (2012). PrimerBank: a PCR primer database for quantitative gene expression analysis, 2012 update. Nucleic Acids Res. 40, D1144–D1149. 10.1093/nar/gkr1013.

65. Xi, H., Fujiwara, W., Gonzalez, K., Jan, M., Liebscher, S., Handel, B., Schenke-Layland, K., and Pyle, A.D. (2017). In Vivo Human Somitogenesis Guides Somite Development from hPSCs. Cell Reports 18, 1573–1585. 10.1016/j.celrep.2017.01.040.

